# The kallikrein-kinin system is falling into pieces: bradykinin fragments are biological active peptides

**DOI:** 10.1101/2020.09.14.296004

**Authors:** Igor Maciel Souza-Silva, Cristiane Amorim de Paula, Anderson Kenedy Santos, Vívian Louise Soares de Oliveira, Isabella Domingos da Rocha, Maísa Mota Antunes, Lídia Pereira Barbosa Cordeiro, Vanessa Pereira Teixeira, Sérgio Ricardo Aluotto Scalzo Júnior, Flávio Almeida Amaral, Jarbas Magalhães Resende, Marco Antônio Peliky Fontes, Gustavo Batista Menezes, Silvia Guatimosim, Robson Augusto Souza Santos, Thiago Verano-Braga

## Abstract

**Background and purpose:** Bradykinin [BK-(1-9)] is an endogenous peptide involved in many physiological and pathological processes, such as cardiovascular homeostasis and inflammation. The central dogma of the kallikrein-kinin system is that BK-(1-9) fragments are biologically inactive. In this manuscript, we proposed to test whether these fragments were indeed inactive.

**Experimental Approach:** Nitric oxide (NO) was quantified in human, mouse and rat cells loaded with DAF-FM after stimulation with BK-(1-9), BK-(1-7), BK-(1-5) and BK-(1-3). We used adult male rat aortic ring preparation to test vascular reactivity mediated by BK-(1-9) fragments. Changes in blood pressure and heart rate was measured in conscious adult male rats by intraarterial catheter method.

**Key results:** BK-(1-9) induced NO production in all cell types tested by B2 receptor activation. BK-(1-7), BK-(1-5) and BK-(1-3) also induced NO production in all tested cell types but this response was independent of the activation of B1 receptor and/or B2 receptor. BK-(1-7), BK-(1-5) or BK-(1-3) induced only vasorelaxant effect and in a concentration-dependent fashion. Vasorelaxant effects for BK-(1-7), BK-(1-5) or BK-(1-3) were independent of the kinin receptors. Different administration routes (*i.e*., intravenous or intra-arterial) did not affect the observed hypotension induced by BK-(1-7), BK-(1-5) or BK-(1-3). Importantly, these observations diverged from the BK-(1-9) results, highlighting that indeed the BK-(1-9) fragments do not seem to act via the classical kinin receptors.

**Conclusions and implications:** In conclusion, BK-(1-7), BK-(1-5) and BK-(1-3) are biologically active components of the kallikrein-kinin system. Importantly, observed pathophysiological outcomes of these peptides are independent of B1R and/or B2R activation.

## 1. INTRODUCTION

Bradykinin [BK-(1-9)] is a peptide-hormone of the kallikrein-kinin system (KKS) which actions were first characterized by an elegant study by Rocha e Silva and coworkers in 1949 (37). The authors observed pronounced hypotension and slow acting contraction of guinea-pig uterus due to administration of plasma treated with snake venom or trypsin. A decade later, effects of purified BK-(1-9) were unveiled by Elliott and coworkers (8). The authors observed that BK-(1-9) induced hypotension, slow contraction of smooth muscle and pro-inflammatory effects. Since then, a considerable amount of knowledge was obtained concerning the KKS, such as: i) inactivation of BK-(1-9) by angiotensin converting enzyme (ACE) (49); ii) elucidation of two proteoforms of kininogens (14); iii) identification of des-Arg^9^-BK-(1-9) [BK-(1-8)] as a biologically active component of KKS (33); iv) cDNA cloning of B2 receptor (B2R), a selective target for BK-(1-9) (24); v) cDNA cloning of B1 receptor (B1R), a selective target for BK-(1-8) (25) and; vi) FDA approval of the selective B2R antagonist Icatibant as the first drug targeting the KKS (1).

BK-(1-9) fragments have been somewhat overlooked for the past 40 years. In the late 60’s, it was reported that BK-(1-9) had a half-life of about 17 seconds in the plasma (10), being metabolized in the pulmonary circulation (38). Given its short half-life, several studies aimed at characterizing the potential activity of BK-(1-9) fragments in different species and using different protocols. However, all studies concluded that the fragments derived from BK-(1-9) were devoid of biological activity, except for BK-(1-8) (30, 32-34, 44). From extensive studies regarding BK-(1-9) proteolysis, it is consensus that BK-(1-8), BK-(1-7) and BK-(1-5) are the major proteolytic fragments, being the latter the most stable fragment in plasma (19, 41). Important to mention that a study reported that BK-(1-5) was biologically active (26) but this information was disregarded by publications reviewing the KKS.

Our research group reported that a cryptic peptide comprising the amino acid sequence Lys-Pro-Pro from the scorpion venom *Tityus serrulatus* (45) acts on the B2R to increase nitric oxide (NO) production and vasorelaxant effect in rat aortic rings (46). Lys-Pro-Pro shares physical-chemical features with the C-terminus of BK-(1-9) (Arg-Pro-Pro) [BK-(1-3)], as both contain a basic residue and two proline residues in tandem. Since Lys-Pro-Pro elicits biological responses, we hypothesized that BK-(1-3) would also be biologically active. Given these circumstances, we proposed in this manuscript a re-evaluation of the biological activity status of BK-(1-7), BK-(1-5) and BK-(1-3). Contrary to current knowledge that BK-(1-9) fragments are inactive, we report that BK-(1-7), BK-(1-5) and BK-(1-3) possess biological activity, assessed in different models. These peptides led to an increase in NO production and vasodilation independently of the kinin receptors activation. We also report *in vivo* actions of BK-(1-9) related peptides.

## 2. METHODS

### 2.1 Reagents

Bradykinin fragments BK-(1-7) [RPPGFSP] and BK-(1-3) [RPP] were purchased from GenOne Biotechnologies. Unless otherwise stated, all reagents were purchased by Sigma-Aldrich.

### 2.2 Peptide synthesis, purification and characterization

BK-(1-9) [RPPGFSPFR], BK-(1-5) [RPPGF] and the B1R selective antagonist Lys-(des-Arg^9^-Leu^8^)-BK-(1-9) [KRPPGFSPL] were synthesized by solid-phase peptide synthesis employing the Fmoc strategy (5), using NovaSyn-TGA resins (Merck). The final unprotected peptides were purified by HPLC using a DiscoveryBIO® Wide Pore C18 semi-preparative column. Purified products were analyzed by MALDI TOF/TOF Autoflex III Smartbeam mass spectrometer (Bruker Daltonics) for synthesis quality control.

### 2.3 Animals

All animal procedures were approved by local animal ethical committee (CEUA-UFMG protocol 213/2016). Adult male Wistar rats of 8-12 weeks old, neonatal male Wistar rats of 3-5 days old and adult male C57Bl/6 mice were obtained by CEBIO-UFMG. B1 and B2 receptor knockout (B1RKO and B2RKO, respectively) male mice were obtained from the animal facility of the Department of Biochemistry and Immunology at UFMG. All animals were housed at the animal facility of the Department of Physiology and Biophysics (UFMG) in accordance with the current edition of the NIH *Guide for the Care and Use of Laboratory Animals*, with chow and water *ad libitum*.

### 2.4 Nitric oxide measurements in cell culture

Human glioblastoma cell line U-87 MG was cultured in DMEM-F12 cell media, supplemented by 10% fetal bovine serum (FBS) (Cultilab) and 1% of antibiotic/antimycotic solution (Thermo-Fisher) at 37°C. Rat neonatal cardiomyocytes were isolated, as previously reported (12). Cells were maintained with DMEM-F12 cell media supplemented by 10% FBS and 1% of antibiotic/antimycotic solution at 37°C until usage. Mice ventricular adult cardiomyocytes were isolated by collagenase type-2 (Worthington), as previously shown (36) and kept in Tyrode media (1.3 x 10^−1^ mol.L^-1^ NaCl; 5.4 x 10^−3^ mol.L^-1^ KCl; 2,5 x 10^−2^ mol.L^-1^ HEPES; 5 x 10^−4^ mol.L^-1^ MgCl_2_; 3.3 x 10^−4^ mol.L^-1^ NaH_2_PO_4_; 1.8 x 10^−3^ mol.L^-1^ mM CaCl_2_; 2.2 x 10^−2^ mol.L^-1^ glucose) until usage.

For NO quantification, cells were starved with Hank’s Balanced Salt Solution (HBSS) for 1 hour and then loaded with DAF-FM diacetate (Thermo-Fisher) at a final concentration of 5 x 10^−6^ mol.L^-1^ for 30 minutes. Cells were rinsed with HBSS and stimulated for 15 minutes with BK-(1-9), BK-(1-7), BK-(1-5) or BK-(1-3), at the final concentration of 10^−7^ mol.L^-1^, diluted with sterile saline. Kinin receptors involvement in NO production by the tested peptides was assessed by selective antagonism of B1 and B2 receptors with Lys-(des-Arg^9^-Leu^8^-BK-(1-9) (B1RB) and HOE-140 (Tocris) (B2RB), respectively. Antagonists were used at a final concentration of 10^−7^ mol.L^-1^, diluted with sterile saline, prior stimulation by BK-(1-9) or its fragments. A concentration-response curve was made by stimulating male neonatal rat cardiac myocytes with BK-(1-9) and its fragments at the final concentration of 10^−7^, 10^−8^ and 10^−9^ mol.L^-1^, diluted with sterile saline, for 15 minutes. After stimulation, cells were fixed with a 4% paraformaldehyde solution for 15 minutes and then mounted. Fluorescence emitted from cells was captured by either a confocal microscope Zeiss LSM 5 LIVE (Zeiss) or a Nikon Eclipse Ti (Nikon), with λ_Ex_ = 495nm and λ_Ex_ = 505nm. Fluorescence emitted by cells was quantified by ImageJ (40).

### 2.5 Vascular reactivity

Vascular reactivity of BK-(1-9) and its fragments were assessed as previously described (35). Briefly, 8-12 weeks old male Wistar rats were euthanized by decapitation. Thoracic aorta was quickly dissected, connective and adipose tissue were removed and then the vessel was cut into 3-5mm long rings. The rings were carefully placed between two stainless-steel stirrups and coupled to an isometric force transducer. The preparation was kept at 37°C, bathed in Krebs-Henseleit salt solution (1.18 x 10^−1^ mol.L^-1^ NaCl; 4.7 x 10^−3^ mol.L^-1^ KCl; 1.17 x 10^−3^ mol.L^-1^ KH_2_PO_4_; 1.12 x 10^−3^ mol.L^-1^ MgSO_4_.7H_2_O; 1.12 x 10^−2^ mol.L^-1^ Glucose; 2.5 x 10^−2^ mol.L^-1^ NaHCO_3_; 2.5 x 10^−3^ mol.L^-1^mM CaCl_2_.2H_2_O, pH fixed at 7.2-7.4). Solution was lightly bubbled with 95% O_2_/5% CO_2_. Endothelial cells of some aortic rings were removed by gently scrubbing the rings with a thin wire to evaluate endothelial role in vasorelaxation. Aortic rings were stabilized for at least 60 minutes under the passive tension of 1.6 g. During the stabilization period, Krebs-Henseleit solution was replaced every 15 minutes and passive tension was corrected to 1.6 g if needed. Endothelium integrity was assessed by pre-contracting the rings with phenylephrine at the final concentration of 10^−7^ mol.L^-1^, diluted with saline. After stabilization of contraction, acetylcholine was added at the final concentration of 10^−6^ mol.L^-1^, diluted with saline, to induce endothelium-dependent vasorelaxation. Endothelium was considered intact when vasorelaxation mediated by acetylcholine was higher than 80%. For experiments without functional endothelium [E (-)], only preparations with no response to acetylcholine were used.

Vasorelaxation mediated by BK-(1-9), BK-(1-7), BK-(1-5) and BK-(1-3) was assessed in endothelium-intact rings by constructing a concentration-response curve for each peptide, after pre-contraction of rings with phenylephrine at a final concentration of 10^−7^ mol.L^-1^, diluted with saline. Tested concentrations ranged from 10^−12^ to 10^−6^ mol.L^-1^, in which all concentrations were diluted with saline. A saline curve was also built by adding the same volume to the media as in the curve with peptides, as a negative control. Endothelium dependency to those effects was assessed in denuded rings (E (-)). Dependency of NO and prostanoids in vasorelaxation was assessed by blocking nitric oxide synthases (NOS) and cyclooxygenases (COX) with N_ω_-nitro-L-arginine methyl ester (L-NAME) and indomethacin, respectively. Both inhibitors were used at the final concentration of 10^−6^ mol.L^-1^, in which L-NAME was diluted with saline and indomethacin was diluted in DMSO. Kinin receptors involvement on BK-(1-9), BK-(1-7), BK-(1-5) and BK-(1-3) induced vasorelaxation was assessed by selectively blocking B1, B2 or either B1 and B2 receptors with Lys-(des-Arg^9^-Leu^8^)-BK-(1-9), HOE-140 or a mixture of both, all at the final concentration of 10^−7^ mol.L^-1^, diluted with saline. Inhibitors and antagonists were incubated in preparations 20 minutes prior contraction by phenylephrine.

### 2.6 Blood pressure measurements in conscious rats

The day before blood pressure measurement, 8-12 weeks old male Wistar rats were anesthetized by ketamine/xylazine (60 mg.kg^-1^ / 9 mg.kg^-1^, respectively, *i.p*). A polyethylene catheter was inserted into the abdominal aorta through the femoral artery for blood pressure measurements and another catheter was inserted into the inferior cava vein through the femoral vein or in the aorta through the left common carotid artery for intravenous (*i.v*.) or intraarterial (*i.a*) administration, respectively, of BK-(1-9) and its fragments. Rats were individually housed and left to recover for 24 hours. On the following day, rats were coupled to a pressure transducer and submitted to a period of stabilization until cardiovascular parameters were stable.

First, a dose-response relationship was built to assess whether BK-(1-9) and its fragments were able to elicit biological responses *in vivo*. In order to do so, 0.1mL of BK-(1-9), BK-(1-7), BK-(1-5) or BK-(1-3) was administered (*i.v*.) *in bolus*, corresponding to a dose of 2.5 x 10^−9^ mol (2.5 nmol), 5 x 10^−9^ mol (5 nmol) or 10^−8^ mol (10 nmol) per animal, in which all doses were diluted in sterile saline. Sterile saline was injected at the same volume as control. To further explore the *in vivo* effect mediated by BK-(1-9) and its related fragments, 0.1 mL of each peptide was administered *in bolus*, corresponding to a dose of 10^−8^ mol (10 nmol per animal, diluted in sterile saline), in the arterial or venal circulation of adult male rats to assess whether injections on different vascular beds would alter the response of these molecules. We also decided to test whether *in vivo* blockage of ACE would have an impact on the observed responses of BK-(1-9), BK-(1-7), BK-(1-5) and BK-(1-3). After administration of captopril (5mg.kg^-1^, *i.v*., diluted in sterile saline) and stabilization of cardiovascular parameters, animals were subjected to the administration 0.1 mL of BK-(1-9) or its related peptides, corresponding to the dose of 10^−8^ mol per animal, diluted in sterile saline.

### 2.7 Relative gene expression by quantitative polymerase chain reaction (qPCR)

Total RNA from U-87 MG, male rat neonatal cardiomyocytes, male adult ventricular mice cardiomyocytes and male rat thoracic aorta was extracted with TRIzol reagent (Thermo-Fisher) and cDNA was synthesized using First Strand cDNA Synthesis (Thermo-Fisher) kit following manufacturer instructions. qPCR was performed in a StepOnePlus Real Time PCR system (Applied Biosystem) using Maxima SYBR PCR 2x (Thermo-Fisher) and primers (Table 1) (Síntese Biotecnológica) at the final concentration of 6 x 10^−7^ mol.L^-1^. The cDNA was diluted in RNAse free water 1:100. Thermal cycling protocol was as follows: i) denaturation at 95°C for 5 minutes followed by; ii) 50 cycles of denaturation at 95°C for 10 seconds; iii) annealing/extension at 60°C for 1 minute. Relative expression of the selected genes was performed by 2^-ΔΔCt^ method (21) using the housekeeper gene S26 as normalizer.

### 2.8 Statistical analysis

All statistical analyses were performed in GraphPad Prism v.7.0 in which p < 0.05 was considered statistically significant. Confocal microscopy, plasma extravasation and nociception experiments were analyzed by One-way ANOVA with Tukey’s multiple comparison as a *post-hoc* test. Vascular reactivity experiments and arterial blood pressure measurements were analyzed by Two-way ANOVA with Tukey’s multiple comparison as a *post-hoc* test. Results are expressed as mean ± SEM.

## 3. RESULTS

### 3.1 BK-(1-9) fragments induce NO production *in vitro*

Bearing the structural resemblance of BK-(1-9) and BK-(1-8) to BK-(1-7), BK-(1-5) and BK-(1-3), notably by the presence of the Arg-Pro-Pro sequence on their N-terminal, which is similar to the Lys-Pro-Pro sequence of a cryptide from the scorpion *Tityus serrulatus* venom that is able to induce NO production via B2 receptor (45, 46), we hypothesized that BK-(1-9) related fragments induced NO production *in vitro* as well. We decided to test our hypothesis with different cell types to evaluate whether NO production induced by BK-(1-9) metabolites are relevant among the most used species in pharmacological studies. As expected, 10^−7^ mol.L^-1^ of BK-(1-9) led to an increase of NO production on immortalized human glioblastoma cells (U-87 MG), neonatal male rat cardiomyocytes and male adult mouse ventricular myocytes. 10^−7^ mol.L^-1^ of BK-(1-7), BK-(1-5) and BK-(1-3) were also able to induce NO production in these cell types (Figure 1).

**Figure 1.**
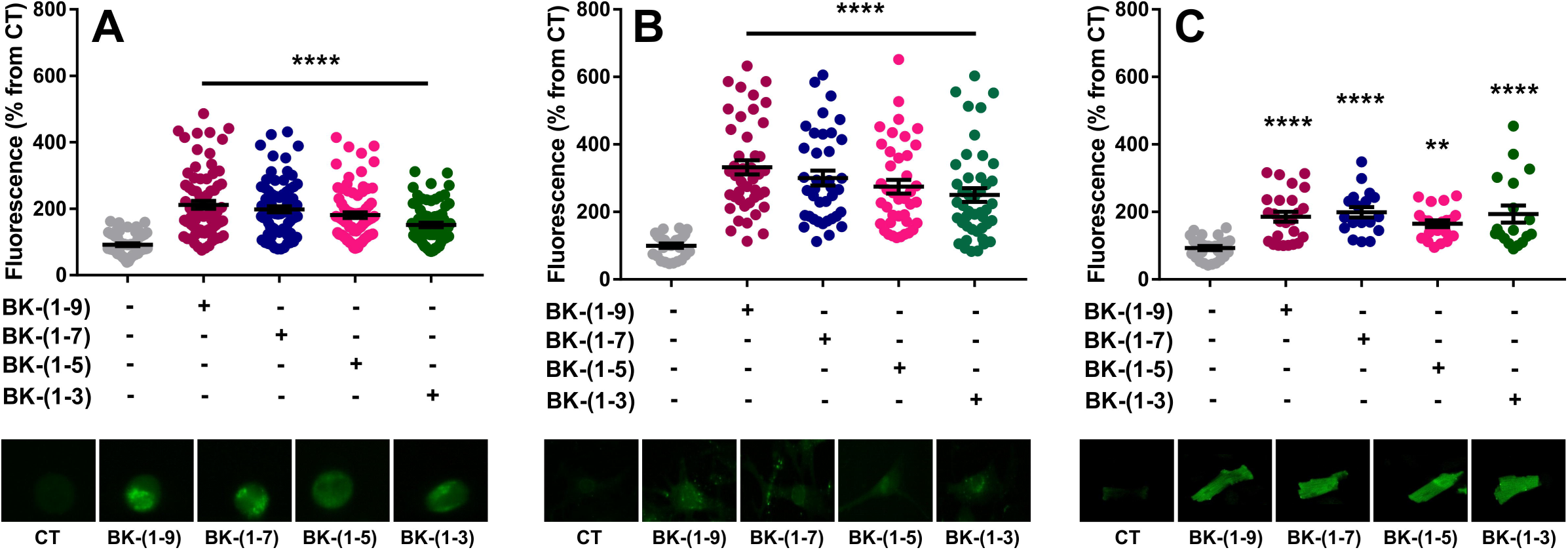
NO production mediated by BK-(1-9) fragments in different cell types. *In vitro* NO production after treating cells with BK-(1-9), BK-(1-7), BK-(1-5), BK-(1-3) at 10^−7^ mol.L^-1^ or sterile saline (control, vehicle for peptide dilution) for 15 minutes. Quantification of NO production in: (A) U-87 MG (n = at least 61 cells/condition); (B) adult male ventricular mice cardiomyocytes, and (n = at least 18 cells/condition) (C) male neonatal rat cardiomyocytes (n = at least 29 cells/condition). One-way ANOVA with Tukey’s multiple comparison *post-hoc* test ** p < 0.01 and **** p < 0.0001 when compared to control. We performed three independent experiments for all data shown. A representative confocal microscopy picture for each experimental condition is shown. Values are expressed as mean ± SEM.

We decided to build a concentration-response curve to analyze the pharmacological characteristics of NO production mediated by BK-(1-9), BK-(1-7), BK-(1-5) and BK-(1-3) in neonatal male rat cardiomyocytes. At all concentrations tested, BK-(1-9) was able to elicit NO production, in which no significant statistical difference was noted among the concentrations tested (Figure 2A). BK-(1-7) was able to induce NO production on neonatal male rat cardiomyocytes only upon simulation at the final concentrations of 10^−7^ and 10^−8^ mol.L^-1^ (Figure 2B). BK-(1-5) (Figure 2C) and BK-(1-3) (Figure 2D) were able to induce NO production even at 10^−9^ mol.L^-1^ but at this tested concentration, a significant lower NO production was detected. These results suggest that BK-(1-7), BK-(1-5) and BK-(1-3) are not only byproducts of BK-(1-9) proteolysis since, at least *in vitro*, we were able to detect NO production in cell lines derived from the most well studied organisms in pharmacology.

**Figure 2.**
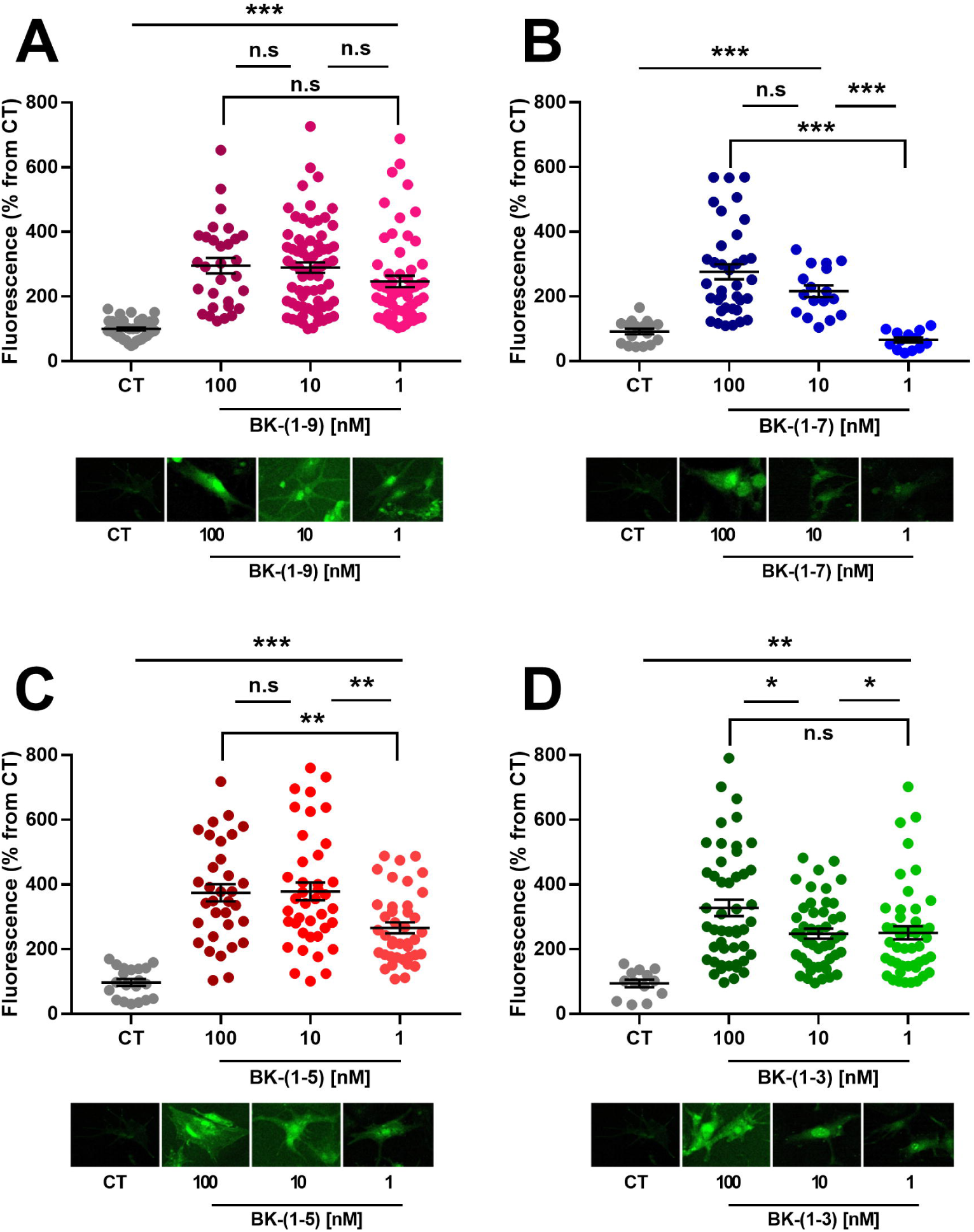
Concentration-response curve of NO production mediated by BK-(1-9) and its fragments on neonatal rat cardiomyocytes. Quantification of NO production in male neonatal rat cardiomyocytes mediated by 100, 10 or 1nM of (A) BK-(1-9); (B) BK-(1-7); (C) BK-(1-3); or (D) BK-(1-3) (n = at least 13 cells/condition). Sterile saline (used for peptide dilution) was employed at the same volume as a control to assess basal NO production from cells. One-way ANOVA with Tukey’s multiple comparison *post-hoc* test * p < 0.05, ** p < 0.01, *** p < 0.001 and n.s. non-significant. We performed three independent experiments for all data shown. A representative confocal microscopy picture for each experimental condition is shown. Values are expressed as mean ± SEM.

### 3.2 *In vitro* NO production induced by BK-(1-7), BK-(1-5) or BK-(1-3) is not mediated by the kinin receptors

To evaluate whether BK-(1-7), BK-(1-5) and BK-(1-3) perform their actions upon activation of kinin receptors *in vitro*, we pre-incubated cells with B1R or B2R selective antagonists, respectively, Lys-des-Arg^9^-BK-(1-9) (B1RB) or HOE-140 (B2RB). In human glioblastoma cell line (U-87 MG), BK-(1-9)-induced NO production was abolished by selectively blocking B2R or by blocking both kinin receptors (Figure 3A). The effects of BK-(1-7) (Figure 3B), BK-(1-5) (Figure 3C) and BK-(1-3) (Figure 3D) were not affected by these antagonists. We observed the same response profile in neonatal rat cardiomyocytes, where BK-(1-9) (Figure 3E) response was abolished by selective antagonism of B2R or by blocking both kinin receptors. No alteration on NO production induced by BK-(1-7) (Figure 3F), BK-(1-5) (Figure 3G) or BK-(1-3) (Figure 3H) was observed. These results suggest that BK-(1-9) fragments do not act upon B1R and/or B2R activation to induce NO production *in vitro*.

**Figure 3.**
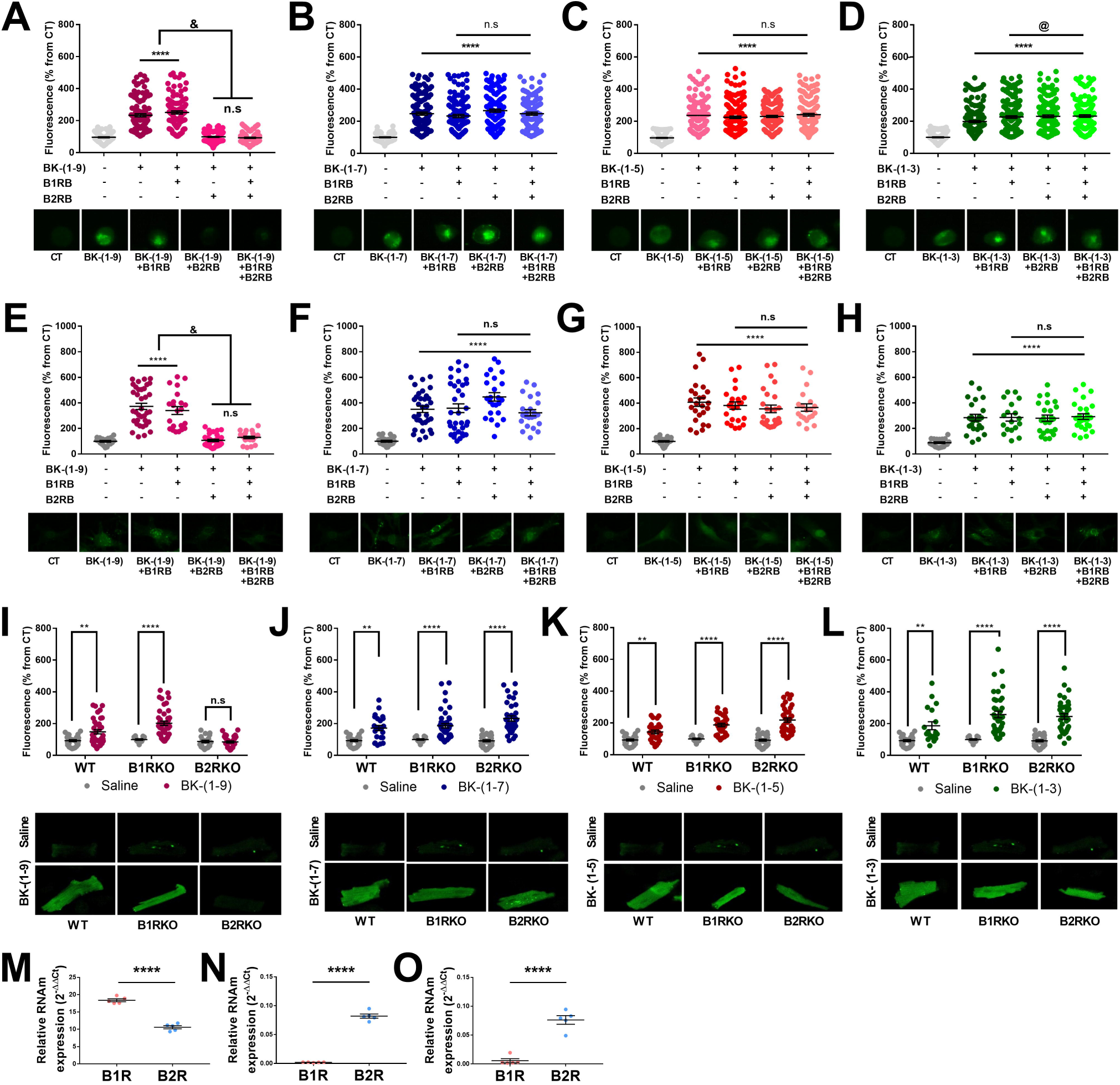
BK-(1-9) fragments do not rely on the activation of kinin receptors to induce NO production *in vitro*. Bradykinin type 1 receptor (B1R) was selectively blocked by Lys-(des-Arg^9^-Leu^8^)-BK-(1-9) (B1RB) and bradykinin type 2 receptor (B2R) was selectively blocked by HOE-140 (B2RB). Both antagonists were used at a final concentration of 10^−7^ mol.L^-1^ for 15 minutes prior to cell stimulation with (A) BK-(1-9) (n = at least 99 cells/condition), (B) BK-(1-7) (n = at least 86 cells/condition), (C) BK-(1-5) (n = at least 126 cells/condition) and (D) BK-(1-3) (n = at least 147 cells/condition) at 10^−7^ mol.L^-1^ or sterile saline for 15 minutes in U-87 MG. NO quantification in cardiomyocytes isolated from male neonatal rat hearts mediated by (E) BK-(1-9) (n = at least 19 cells/condition), (F) BK-(1-7) (n = at least 21 cells/condition), (G) BK-(1-5) (n = at least 20 cells/condition) and (F) BK-(1-3) (n = at least 18 cells/condition). NO quantification in cardiomyocytes isolated from knockout mice for either B1R or B2R, or wild-type (WT) following (I) BK-(1-9) (n = at least 15 cells/condition); (J) BK-(1-7) (n = at least 15 cells/condition); (K) BK-(1-5) (n = at least 15 cells/condition) or (L) BK-(1-3) (n = at least 15 cells/condition) stimulation (always using 10^−7^ mol.L^-1^ of peptide for 15 minutes). Sterile saline (used as diluent for peptides and antagonists) was used as control for NO quantification in human glioblastoma-like cell line and in cardiomyocytes derived from male neonatal rats and male adult mice. We performed three independent experiments for all data shown. A representative confocal microscopy image for each experimental condition is shown. One-way ANOVA with Tukey’s multiple comparison *post-hoc* test. ** p < 0.01, **** p < 0.0001, n.s non-significant when compared to control. @ p < 0.05 when compared to BK-(1-3). Relative mRNA expression of B1R and B2R in (M) U-87 MG, (N) neonatal rat cardiomyocytes, and (O) adult ventricular mouse cardiomyocytes (n = 5). Unpaired t-test; **** p < 0.0001 compared to B1R expression. All results are expressed as mean ± SEM.

Knockout male mice for either B1R (B1RKO) or B2R (B2RKO) were employed to further evaluate the role of kinin receptors on NO production. BK-(1-9) induced NO production only in ventricular cardiomyocytes of B1RKO (Figure 3I). On the other hand, BK-(1-7) (Figure 3J), BK-(1-5) (Figure 3K) or BK-(1-3) (Figure 3L) induced NO production in ventricular cardiomyocytes isolated from B1RKO or B2RKO mice. These effects were comparable to the observed for cardiomyocytes isolated from wild-type (WT) mice.

We used qPCR to quantify the expressions levels for B1R and B2R mRNA on the cell types used in this manuscript. As shown in the Figure 3M, both mRNA receptors are highly expressed in U-87 MG, although expression level of B1R mRNA is at least twice as higher than B2R mRNA. B2R mRNA expression was also detected in neonatal male rat cardiomyocytes (Figure 3N) and male adult mouse ventricular myocytes (Figure 3O). We could not detect the expression of B1R mRNA on these cardiac myocytes. Taken together, these results indicate that BK-(1-7), BK-(1-5) and BK-(1-3) do not act via B1R or B2R to promote NO production *in vitro*, regardless their structural similarity to BK-(1-9) and BK-(1-8).

### 3.3 *Ex vivo* vascular effects of the BK-(1-9) fragments

BK-(1-9) is known to induce vasodilation due to NO and prostacyclin production in endothelial cells (22). Given the fact that BK-(1-7), BK-(1-5) and BK-(1-3) were able to induce NO production *in vitro*, we speculated that these fragments could also promote vasodilation. In male rat aortic rings, BK-(1-9) induced a concentration-dependent vasorelaxation at lower concentrations and vasoconstriction at higher concentrations (Figure 4A). BK-(1-7) (Figure 4B), BK-(1-5) (Figure 4C) and BK-(1-3) (Figure 4D) induced a concentration-dependent vasorelaxation. The percentage of maximum vasorelaxation (E_max_) was 15.09 ± 2.08% for BK-(1-9), 19.36 ± 2.06% for BK-(1-7), 18.45 ± 2.53% for BK-(1-5), and 25.52 ± 5.03% for BK-(1-3).

**Figure 4.**
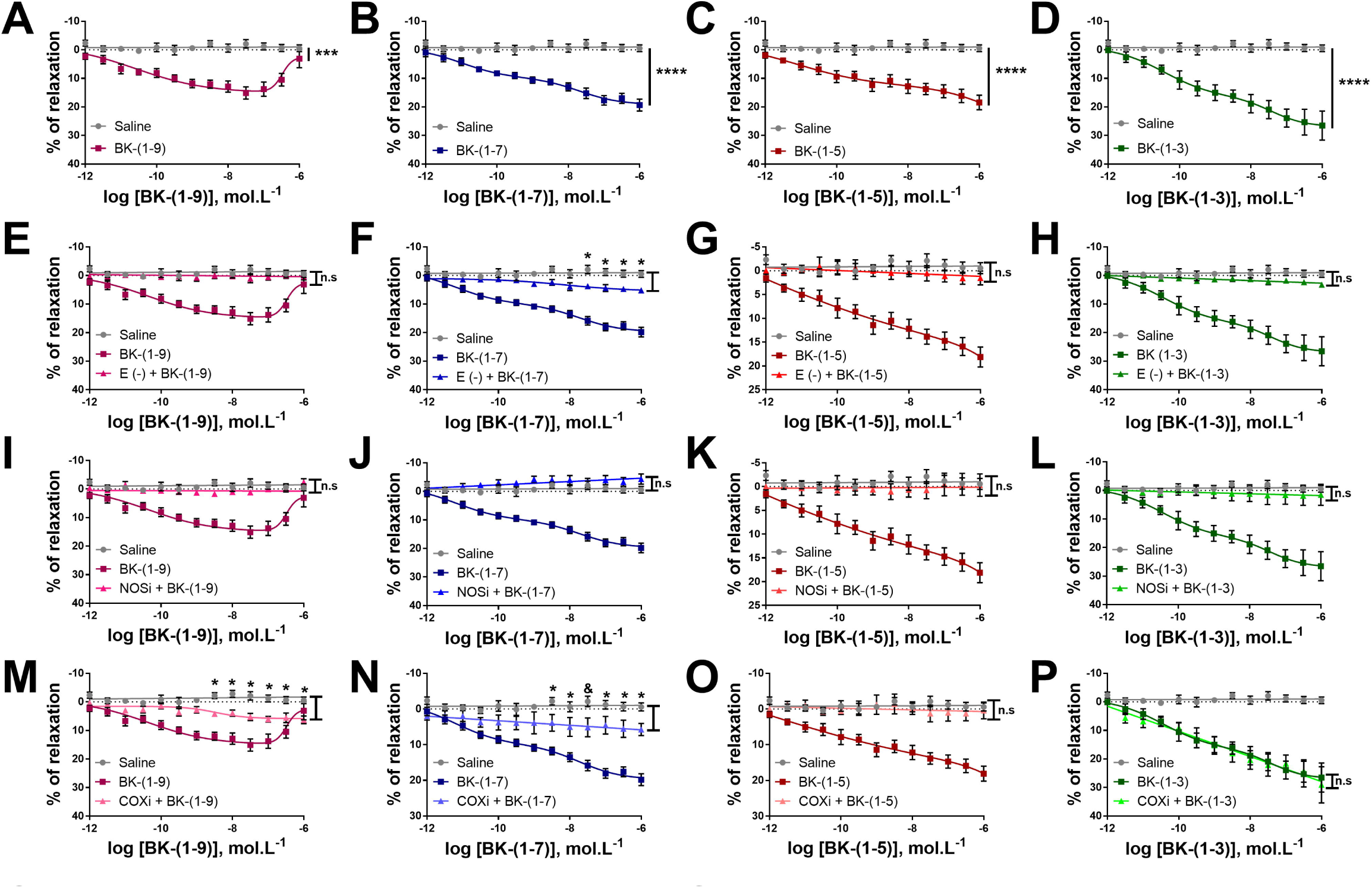
Vascular effects of BK-(1-9) fragments on *ex vivo* preparations of rat aortic rings. Concentration-response curves for (A) BK-(1-9), (B) BK-(1-7), (C) BK-(1-5) and (D) BK-(1-3) performed in aortic rings derived from adult male rats. Removal of functional endothelium (E-) abolished the vasorelaxant effect mediated by (E) BK-(1-9), (G) BK-(1-5) and (H) BK-(1-3) while the effect of (F) BK-(1-7) was partially inhibited. Blockage of NO production by L-NAME at 10^−6^ mol.L^-1^ blunted the effects mediated by (I) BK-(1-9), (J) BK-(1-7), (K) BK-(1-5) and (L) BK-(1-3). The effects of (M) BK-(1-9) and (N) BK-(1-7) were partially blocked by inhibiting COX with indomethacin (diluted with dimethyl sulfoxide) at 10^−6^ mol.L^-1^, while the effect observed for (O) BK-(1-5) was abolished. On the other hand, the effect of BK-(1-3) was not altered when blocking prostanoid production with indomethacin. A control curve employing saline (used for dilution of drugs and peptides, except when mentioned) was built to exclude the possibility of spontaneous vasoactive responses. Two-way ANOVA with Tukey’s multiple compassion *post-hoc* test. * p < 0.05, *** p < 0.001, **** p < 0.0001 and n.s non-significant (n = at least 5 aortic rings from different animals/condition). Results are shown as mean ± SEM., COXi (+ indomethacin), NOSi (+ L-NAME).

Furthermore, the vascular effect of BK-(1-9) was completely blocked when endothelial cells were mechanically removed (Figure 4E) or when NOS were pharmacologically blocked by L-NAME (Figure 4I), whereas this effect was only partially blocked by pharmacological inhibition of cyclooxygenases by indomethacin (Figure 4M). BK-(1-7)-induced vasodilation was partially dependent on endothelial integrity (Figure 4F) and prostanoid production (Figure 4N), while NO production was essential for vasorelaxation (Figure 4J). To promote vasorelaxant effects, BK-(1-5) relied on functional endothelium (Figure 4G), and production of both NO (Figure 4K) and prostanoid (Figure 4O). BK-(1-3)-induced vasorelaxation was only mediated in the presence of functional endothelium (Figure 4H) and NO production (Figure 4L), while prostanoid production (Figure 4P) seems not to play a role on the vasorelaxant effect of this peptide.

### 3.4 The vascular effects of BK-(1-9) fragments are not mediated by the activation of the kinin receptors

Pharmacological blockage of B1R by Lys-(des-Arg^9^-Leu^8^)-BK-(1-9) at 10^−7^ mol.L^-1^ (B1RB), B2R by HOE-140 at 10^−7^ mol.L^-1^ (B2RB) or both receptors by an equimolar mixture B1RB and B2RB at 10^−7^ mol.L^-1^ was performed in male rat aortic rings to assess the relevance of the kinin receptors on vasorelaxation induced by BK-(1-9) fragments. The vasorelaxant effect induced by BK-(1-9) was abolished by B2RB (Figure 5A), B1RB (Figure 5E), and by a mixture of B1RB and B2RB (Figure 5I). Interestingly, vasorelaxation induced by BK-(1-7) (Figures 5B, F and J), BK-(1-5) (Figures 5C, G and K) and BK-(1-3) (Figures 5D, H and L) were not altered by blocking the kinin receptors.

**Figure 5.**
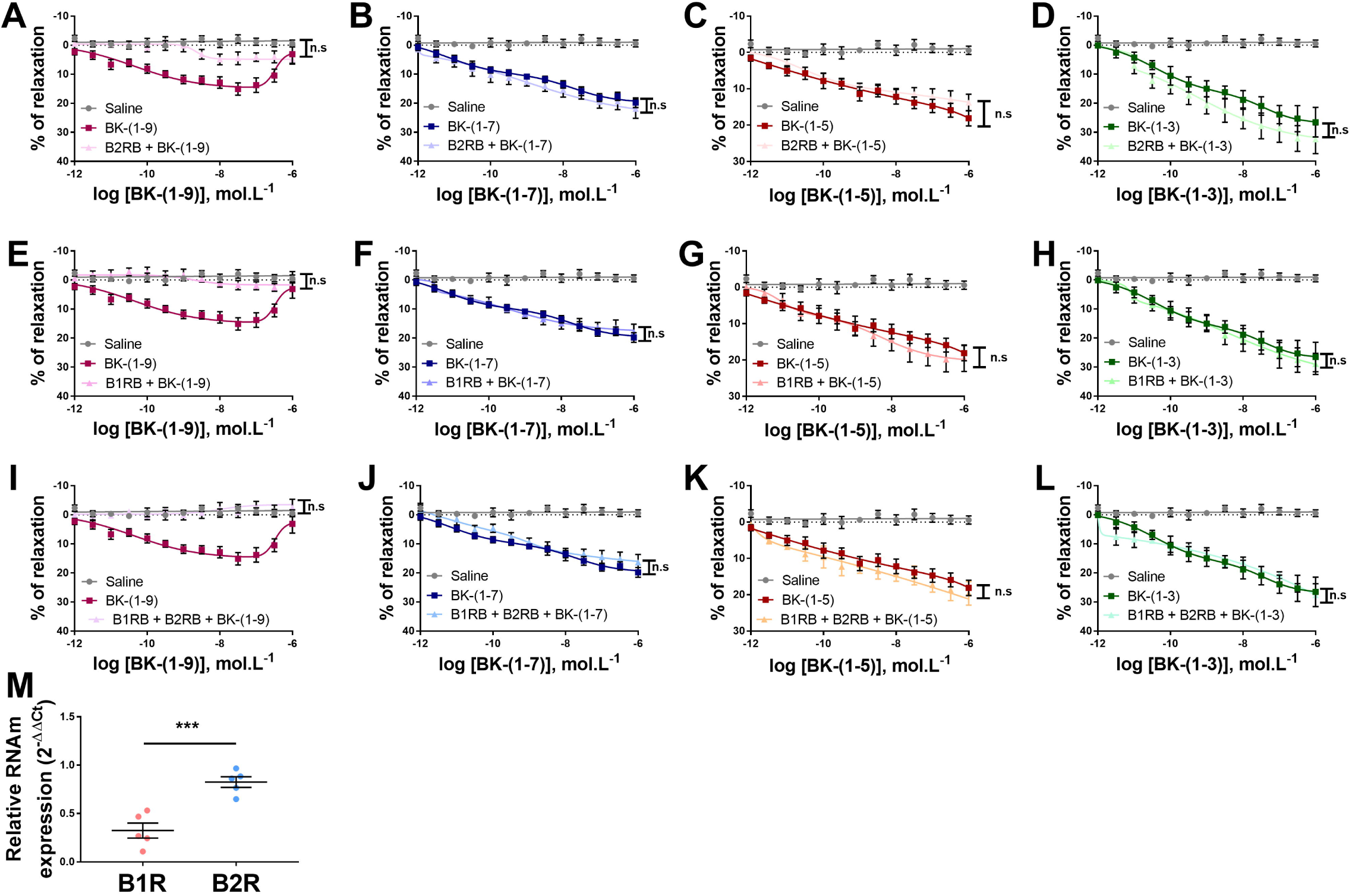
Vasorelaxant effects of BK-(1-9) fragments are independent of the activation of the kinin receptors. Concentration-response curves performed in aortic rings derived from adult male rats. BK-(1-9) response was abolished by blocking (A) B2R with HOE-140 (B2RB) at 10^−7^ mol.L^-1^, (E) B1R with Lys-(des-Arg^9^-Leu^8^)-BK-(1-9) (B1RB) at 10^−7^ mol.L^-1^ or by (I) an equimolar mixture of B1RB and B2RB at 10^−7^ mol.L^-1^ each. BK-(1-7) vasorelaxant response was not altered by (B) B2RB, (F) B1RB or (J) B1RB + B2RB. Response of BK-(1-5) was also not modified by (C) B2RB, (G) B1RB or (K) B1RB + B2RB. For BK-(1-3), selective antagonism of (D) B2R, (H) B1R or (L) both receptors did not alter vasorelaxation. A control curve employing saline (used for dilution of drugs and peptides) was built to exclude the possibility of spontaneous vasoactive responses. Two-way ANOVA with Tukey’s multiple compassion *post-hoc* test; n.s non-significant (n=5). (M) Relative mRNA expression of B1R and B2R on adult male rat thoracic aorta (n = at least 5 aortic rings from different animals/condition). Unpaired t-test. *** p < 0.001 when compared to B1R mRNA expression. All results are shown as mean ± SEM.

Expression of the kinin receptors mRNAs were assessed in rat thoracic aorta by qPCR. Both B1R and B2R mRNA were detected in the rat thoracic aorta, but the expression of B2R mRNA was significantly higher than B1R mRNA (Figure 5M). Taken together, the results indicate that vasorelaxation induced by BK-(1-7), BK-(1-5) and BK-(1-3) are independent of B1R and/or B2R activation.

### 3.5 *In vivo* effects of the BK-(1-9) fragments

Vasodilation is often translated *in vivo* to hypotension due to reduced vascular peripheral resistance. Thus, we aimed at assessing whether BK-(1-7), BK-(1-5) and BK-(1-3) would also promote this physiological outcome. When administered intravenously (i.v) in awaken non-anesthetized male rats, BK-(1-9) induced a transient dose-dependent hypotension followed by a transient increase in heart rate (tachycardia) due to baroreflex activation. BK-(1-7), BK-(1-5) or BK-(1-3) induced a significant hypotension followed by tachycardia but, unlike BK-(1-9), this effect was dose-independent (Figures 6A and B, traces on Supplementary Figure S5). The dose-independent relationship observed for BK-(1-9) fragments could be due to extensive proteolysis of these peptides in the circulation, mainly in the pulmonary circulation. To further explore this, we investigated whether administration of BK-(1-9) and its fragments on a different vascular bed could change the observed responses. Intra-arterial (i.a.) administration of BK-(1-9) led to increased hypotension when compared to i.v. administration (Figures 6C and D, traces on Supplementary Figure S6), most likely because BK-(1-9) is subjected to extensive proteolysis in the pulmonary circulation (39) when i.v. administrated. On the other hand, we could not detect any significant difference when the BK-(1-9) fragments were i.a. administered (Figures 6C and D), suggesting that the BK-(1-9) fragments are somehow more resistant to proteolysis in the pulmonary circulation. Since ACE is the main peptidase involved in BK-(1-9) proteolysis and this enzyme is particularly active in the pulmonary capillaries (29), we decided to perform an *in vivo* inhibition of ACE with captopril to evaluate potential impacts in the cardiovascular effects of the BK-(1-9) fragments. As expected, ACEi led to an exacerbated response of BK-(1-9) (Figures 6E and F, traces on Supplementary Figure S7). On the other hand, the cardiovascular effects of the BK-(1-9) fragments were not affected by ACEi (Figures 6E and F, traces on the Supplementary Figure S7). Taken together, our results suggest that BK-(1-9) proteolytic fragments have an *in vivo* activity. The effects observed for each peptide are most likely of their own since administration on other vascular bed and ACE blockage were not able to alter the responses. We acknowledge that other carboxypeptidases may take part on the metabolization on BK-(1-7), BK-(1-5) and BK-(1-3) to smaller fragments and further investigation is needed to confirm on whether the observed effects of these molecules are of their own.

**Figure 6.**
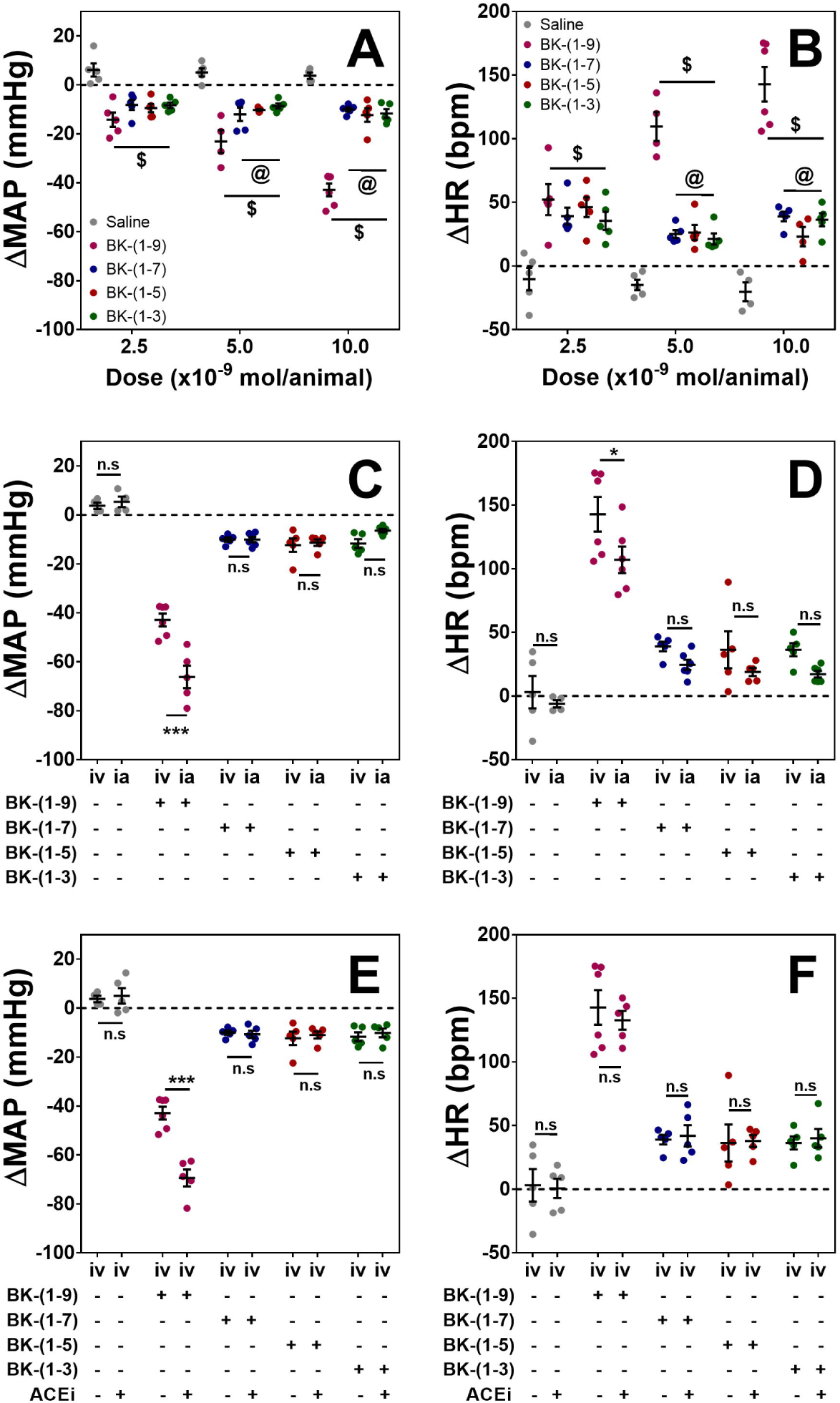
Assessing the involvement of BK-(1-9) fragments on arterial blood pressure of non-anesthetized adult male rats. On (A) and (B) changes of mean arterial pressure (MAP) and heart rate (HR), respectively, mediated by an intravenously (*i.v*.) in bolus injection of BK-(1-9), BK-(1-7), BK-(1-5) or BK-(1-3) at three different doses (2.5 x 10^−9^ mol, 5 x 10^−9^ mol or 10^−8^ mol per animal, 0.1mL). Sterile saline (used for peptide dilution) was administered at the same volume as control. Two-way ANOVA with Tukey’s multiple compassion *post-hoc* test. $ p < 0.0001 when compared to saline. @ p < 0.05 when compared to BK-(1-9) or its fragments (n = 5). On (C) and (D) changes of mean arterial pressure (MAP) and heart rate (HR), respectively, mediated by an intravenously (*i.v*.) or an intra-arterial (*i.a*.) administration of 10^−8^ mol (per animal, 0.1mL) of BK-(1-9) or its fragments. Sterile saline (used for peptide dilution) was administered at the same volume as control. Two-way ANOVA with Sidak’s multiple compassion *post-hoc* test. *** p < 0.001, * p < 0.05, n.s. non-significant (n = 5). On (E) and (F) changes of mean arterial pressure (MAP) and heart rate (HR), respectively, mediated by BK-(1-9) and its fragments with or without pre-treatment with captopril (ACEi) (5mg.kg^-1^ per animal, 0.1mL). Sterile saline (used for peptide dilution) was administered at the same volume as control. Two-way ANOVA with Sidak’s multiple compassion *post-hoc* test. *** p < 0.001, n.s. non-significant (n = 5). All results are shown as mean ± SEM. ACEi (+ captopril).

Apart from its cardiovascular effects, BK-(1-9) is a potent pro-inflammatory agent (4). Thus, we evaluated whether BK-(1-9) fragments would impact nociceptive reflexes and vascular permeability, which are known inflammatory signs. We observed signs of nociceptive reflexes when BK-(1-9) and its fragments were administered *in bolus* either i.v. or i.a. during our *in vivo* assays. Thus, we evaluated nociceptive reflexes in adult male mice and it was possible to observe that BK-(1-7), BK-(1-5) or BK-(1-3) evoked nociceptive reflexes but at significant lower degree of response when compared to BK-(1-9) (Supplementary Figure S8A). Quantification of plasma extravasation in the mice footpads induced by BK-(1-9), BK-(1-7), BK-(1-5) or BK-(1-3) was made possible by the quantification of Evans’ Blue in the peptide administration site. BK-(1-9) was able to increase vascular permeability in mice footpads while no significant effect was observed for BK-(1-7), BK-(1-5) or BK-(1-3) (Supplementary Figure S8B).

## 4. DISCUSSION

Since the characterization of BK-(1-9) activity in 1949 (37), many groups aimed to characterize the potential biological effects of its plasma fragments. However, research made on this regard built the KKS central dogma that BK-(1-9) fragments are biologically inactive (30, 32, 34, 44). This scenario was partially dismissed when Regoli *et al*. (33) characterized the activity of BK-(1-8) and postulated the existence of two kinin receptors, which was later confirmed by the cloning of B2R (24) and B1R (25). Reports on another bioactive fragment of BK-(1-9) emerged almost two decades later when Hasan *et al*. (15) demonstrated that BK-(1-5) inhibited alpha-thrombin-induced platelet aggregation and secretion. Later, Morinelli *et al*. (26) reported that BK-(1-5) increased the survival rate of rats in a sepsis model. However, the scientific community seems to have disregarded these findings and the current knowledge that still prevails is that BK-(1-9) fragments are biologically inactive products of BK-(1-9) plasma proteolysis. In this manuscript, we demonstrated biological activity of BK-(1-9) fragments in different species, upgrading their biological status on KKS.

We reported a significant activity of BK-(1-7), BK-(1-5) and BK-(1-3) on NO production *in vitro* using male neonatal rat and male adult mouse cardiomyocytes as well as in human glioblastoma cells. The usage of cardiomyocytes aligns with BK-(1-9) importance in heart physiology, since it is well known that the peptide can induce NO production in cardiac myocytes (16). We decided to use an immortalized glioblastoma-like cell line derived from a human patient (U-87 MG) since this cell line highly expresses mRNA for both B1 and B2 receptors (3), which most likely translates to high density of these receptors in the cell membrane. One may argue that the concentration used to stimulate cells was fairly high and not physiologically possible, but the usage of an high attack dose to stimulate cells is frequently used in pharmacological studies regarding Ang-(1-7) (6), alamandine (7) and BK-(1-9) *in vitro* actions (16). We acknowledge that a concentration-response curve is needed to confirm that BK-(1-9) fragments action on inducing NO production is not a non-selective event and we were able to show in male rat neonatal cardiomyocytes that BK-(1-9), BK-(1-5) and BK-(1-3) displayed activity at the lowest concentration used (10^−9^ mol.L^-1^), whereas BK-(1-7) was only active at the higher concentrations (10^−7^ and 10^−8^ mol.L^-1^). Taken together, the *in vitro* results suggest BK-(1-9) fragments are active, in which the effects are most likely driven by receptor activation and not by a non-specific interaction, given the nanomolar activity of BK-(1-7), BK-(1-5) and BK-(1-3). Further studies are necessary for identifying which target(s) these molecules act upon.

Bearing the structural resemblance between BK-(1-7), BK-(1-5) and BK-(1-3) to BK-(1-9) N-terminus, it was not unreasonable to hypothesize that these fragments act via kinin receptors to elicit the biological outcomes reported in this manuscript. Indeed, structure-activity assays have demonstrated that one of the requisites for a peptide to activate the kinin receptors relies on a basic amino acid (e.g. Arg or Lys) at peptide N-terminus (17). The hypothesis that BK-(1-9) related peptides could activate the kinin receptors was strengthened by the fact that a cryptic scorpion peptide (Lys-Pro-Pro), similar to BK-(1-3), induced NO production in mice cardiomyocytes by the activation of B2R (45, 46). However, to our surprise, none of the observed effects for BK-(1-7), BK-(1-5) and BK-(1-3) were antagonized by B1R and/or B2R antagonists. To rule out the possibility that the BK-(1-9) fragments activate the kinin receptors, we used cardiomyocytes isolated from B1RKO or B2RKO mice, but these fragments were still able to induce NO production. Therefore, we are quite confident to state that BK-(1-7), BK-(1-5) or BK-(1-3) does elicit neither NO production nor vasorelaxation via activation of B1R and/or B2R.

A possible explanation for the observed results lies on the interaction between the peptide C-terminus and kinin receptors. Peptide backbone plays a major role in the activation of kinin receptors, since certain conformations are necessary for proper positioning of peptide side chains into the receptors binding pockets (17). Naturally, reducing the sequence of BK-(1-9) may disrupt the optimal orientation of essential groups responsible for receptor activation. In addition, there are essential amino acids responsible for B1R and B2R selectivity. For example, Phe^8^ of BK-(1-8) is essential for B1R activation, since its removal or substitution for an aliphatic amino acid (*e.g*., Ile) renders loss of activity towards this receptor or a transition of agonist to antagonist activity, respectively (20, 33). Together with proper conformation, Arg^9^ on BK-(1-9) is essential for B2R activation as removal of this residue leads to sharp affinity decrease (17). The structural-activity data corroborates the results herein presented that BK-(1-7), BK-(1-5) and BK-(1-3) do not act upon activation of B1R or B2R and strongly suggests the existence of other target receptor for these BK-(1-9) fragments. Thus, it is imperative that more studies are performed for elucidation of the molecular target(s) for each of these new functional components of KKS.

Blood pressure is controlled by complex mechanisms that modulates cardiac output and peripheral vascular resistance (to review this topic, see (13)). The hallmark effect of BK-(1-9) when administered *in vivo* is a marked hypotension, which was first observed by Rocha e Silva and coworkers in 1949 (37). The hypotensive effect of BK-(1-9) was later attributed to be mainly caused by vasodilation of systemic vessels, leading to a consequent reduction of peripheral vascular resistance (20). We report in this manuscript that BK-(1-7), BK-(1-5) and BK-(1-3) induced a concentration-dependent vasorelaxant effect in aortic rings from male adult rats. However, the molecular mechanisms underlying these effects seem to be different among them. Nonetheless, vasorelaxation induced by BK-(1-9) fragments were translated *in vivo* to transient dose-independent hypotension in awaken rats.

When BK-(1-9) or its fragments were administered either i.v. or i.a. on awaken male rats, we observed augmented locomotion, a nociception sign. It is known that KKS plays a major role in inflammation, as already reviewed (20, 22). To assess whether BK-(1-7), BK-(1-5) and BK-(1-3) played any part in inflammatory events similarly to BK-(1-9), we evaluated the most notable pathophysiological effects by the later peptide: nociception (4) and vascular permeability (18). We observed that BK-(1-7), BK-(1-5) and BK-(1-3) increased nociceptive reflexes on C57Bl/6 mice, but this effect was significant lower compared to BK-(1-9). On the other hand, we did not observe increased plasma extravasation mediated by BK-(1-9) fragments. Even though these results are only preliminary, our data suggest that BK-(1-7), BK-(1-5) and BK-(1-3) are less pro-inflammatory agents than BK-(1-9) and that these molecules may have important outcomes beyond the cardiovascular system, but further experiments are needed to evaluate their potential role in inflammation.

Finally, one could argue that BK-(1-9) fragments are not endogenous peptides. However, we would like to stress that BK-(1-8), BK-(1-7) and BK-(1-5) have been identified as the major BK-(1-9) proteolytic fragments (2, 19, 23, 27, 31, 41). Given the natural occurrence of these peptides and their biological activity reported in this manuscript, it is likely that BK-(1-7) and BK-(1-5) play important roles in physiological and pathological conditions, which requires extensive research to unravel their particularities. Although BK-(1-3) (368.43 Da) was not detected in these studies, its physiological relevancy should not be ruled out since it is challenging to detect peptides with low molecular weight due to technical difficulties. Detection of BK-(1-3) on biological matrixes has proven to be an exceedingly difficult task. While investigating BK-(1-9) plasma metabolism, Sheik and coworkers (42) were able to identify BK-(1-3) in plasma after 3 hours of incubation. Even though a few years later Shima and coworkers (43) described that BK-(1-5) is the most stable BK-(1-9) fragment on plasma, it was proven that this peptide could be further processed into smaller fragments, such as Phe and Gly-Phe. As the authors stated, BK-(1-3) could not be detected in the study due to its rapid metabolization to the amino acids Arg and Pro. In line with this finding, Marshall and coworkers (23) were able to detect Phe after 2 hours of bromo-BK-(1-9) with mass spectrometry techniques, which is a sign for the metabolization of BK-(1-5) and even maybe to BK-(1-3) production. It is worth mentioning that in this protocol the incubation period was relatively short, and this could be the reason why the authors were not able to detect BK-(1-3) on human or rat plasma. Given these circumstances, it is rather unlikely that BK-(1-3) is not produced during BK-(1-9) breakdown but as for now, efforts should be made to develop techniques to identify this peptide in biological matrixes.

## 5. CONCLUSION

Our study shows that the BK-(1-9) proteolytic fragments, BK-(1-7), BK-(1-5) and BK-(1-3), have significant biological activity in human, rat and mouse cell lines. The cardiovascular response of these fragments seems to be in parallel with BK-(1-9). These peptides might have potential biotechnological purpose as we observed significant lower pro-inflammatory responses elicited by them compared to BK-(1-9), but this feature is yet to be further investigated. Apart from being a breakthrough in KKS dogma, the observed biological activity for BK-(1-7), BK-(1-5) and BK-(1-3) open new doors to further characterize this important and yet overlooked system. We believe that KKS is far more complex than previously anticipated, formed by a network of peptides and receptors (yet to be identified) to maintain homeostasis of fine-tuning processes like arterial blood pressure, inflammation, and others.

## ACKNOWLEGMENTS

We are thankful for the technical support of Adriana Campezatto Raabe from the “Laboratório Multiusuários de Proteômica” (LMProt) - “Centro de Laboratórios Multiusuários do Instituto de Ciências Biológicas” (CELAM) at the “Universidade Federal de Minas Gerais” and Jamil Silvano de Oliveira from the “Departamento de Bioquímica e Imunologia” at the “Universidade Federal de Minas Gerais”. We also acknowledge the “Centro de Aquisição e Processamento de Imagens” (CAPI-UFMG) for confocal microscopy experiments.

## STATEMENT OF CONTRIBUTION

I.M.S.S.: Experimental design, peptide synthesis, confocal cell imaging, vascular reactivity, manuscript preparation and revision.

V.P.T., S.R.A.S.J.: Isolation of cardiac myocytes.

A.K.S.: qPCR analysis.

V.L.S.O.: Data acquisition on inflammatory and nociceptive *in vivo* experiments.

I.D.R., C.A.P: *In vivo* arterial blood pressure experiments.

M.M.A: Data acquisition on confocal cell imaging. L.P.B.C.: Peptide synthesis.

F.A.A., J.M.R., G.B.M., M.A.P.F., S.G. and R.A.S.: Experimental design, results discussion and revision of the manuscript.

T.V.B.: Experimental design, results discussion, revision and manuscript submission.

## SUPPORTING INFORMATION

Private sharing link for Figshare data: https://figshare.com/s/463b5aa302cb27e6f153

