## Supplemental Information for "The kallikrein-kinin system is falling into pieces: bradykinin fragments are biological active peptides"

[1]Departamento de Fisiologia e Biofísica, [2]Departamento de Bioquímica e Imunologia, [3]Departamento de Morfologia, [4]Departamento de Química, Universidade Federal de Minas Gerais, Av. Antonio Carlos 6627, Belo Horizonte 31270-901, Brazil.

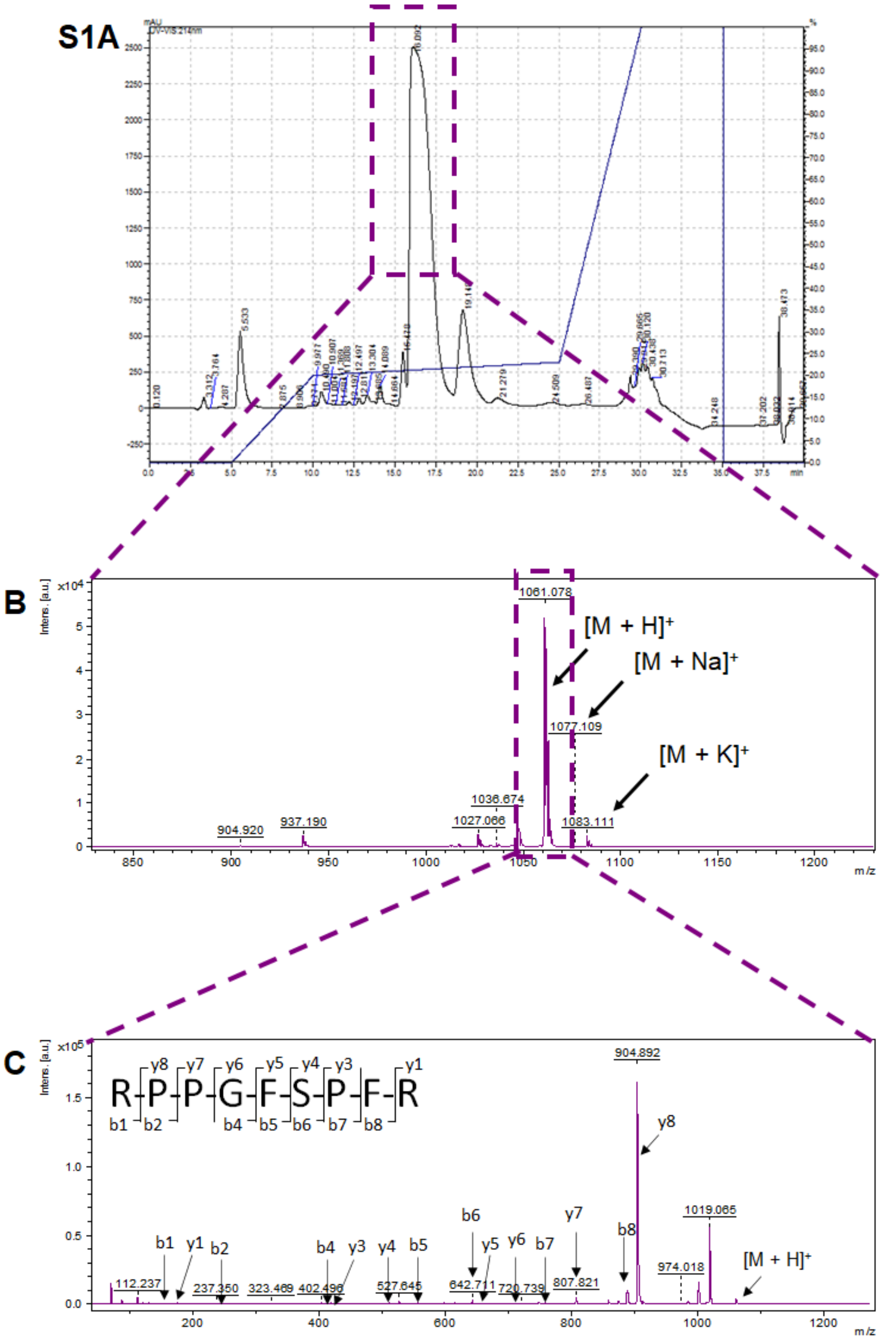


**Supplementary Figure S1 – Purification and characterization of BK-(1-9) synthesis products.** (A) Representative chromatogram from the purification process of crude BK-(1-9). Since the main peak at 16.09 minutes (in the purple dashed box) was not totally resolved, only the superior half of the peak was collected to avoid contamination. (B) MALDI TOF/TOF mass spectrum of the collected fraction in the chromatogram in (A). A prominent ion in 1061.078 $[M + H]^+$, with an experimental mass of 1060.070 Da, corresponded to the theorical mass of BK-(1-9), 1060.56 Da. Adduct ions of $Na^+$ and $K^+$ were also observed, with m/z of 1077.109 and 1083.111, respectively. Even though a few m/z lower than 1061.078 were observed in the spectrum, they were not as intense. In order to confirm the identity of the ion at 1061.078, it was further fragmented resulting in the spectrum at (C). Fragmentation pattern of the ion at 1061.078 confirmed the identity of BK-(1-9).

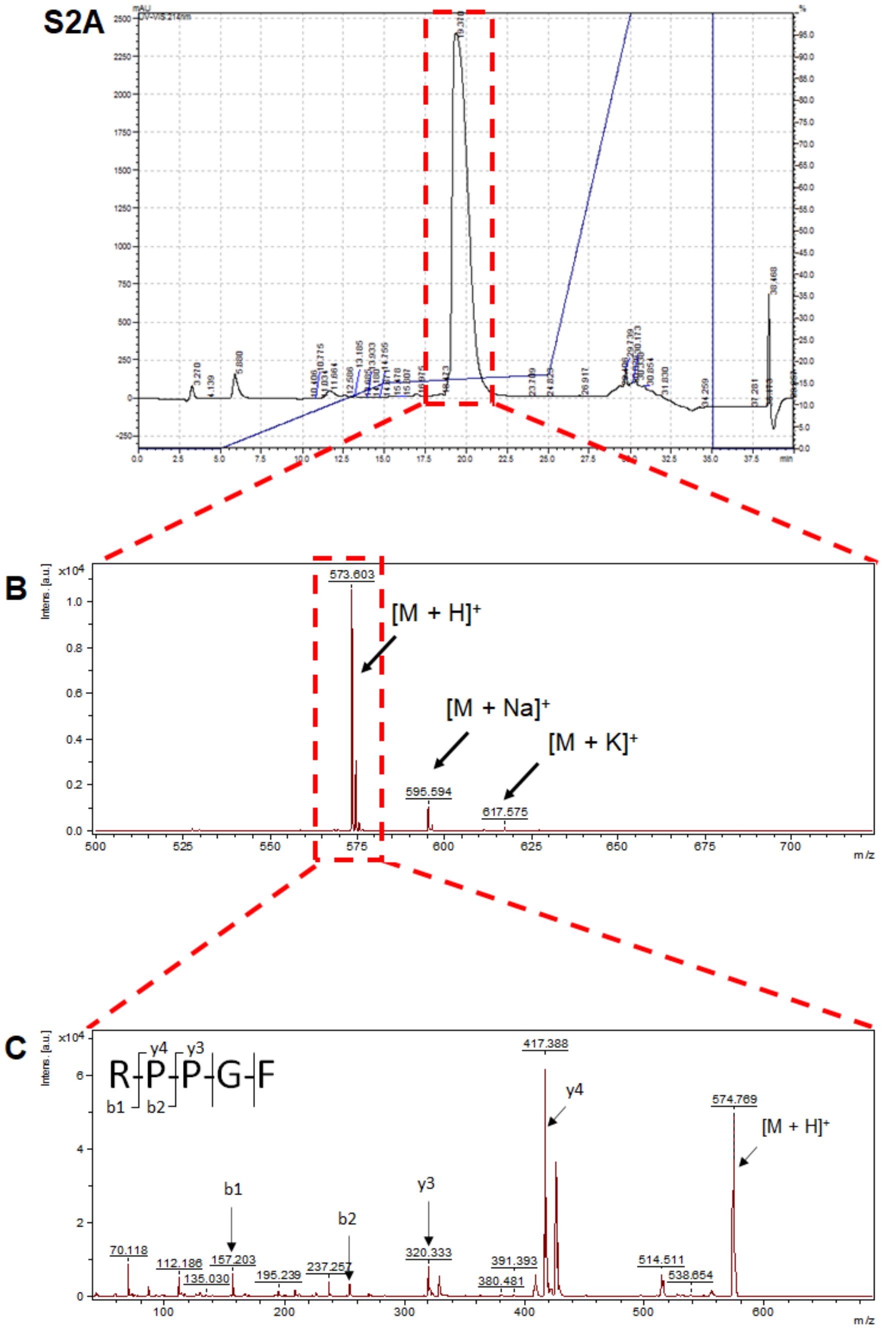


**Supplementary Figure S2 – Purification and characterization of BK-(1-5) synthesis products.** (A) Representative chromatogram from the purification process of crude BK-(1-5). The main peak at 19.37 minutes (in the red dashed box) was collected. (B) MALDI TOF/TOF mass spectrum of the collected fraction in the chromatogram in (A). A prominent ion in 573.603 $[M + H]^+$, with an experimental mass of 572.595 Da, corresponded to the theorical mass of BK-(1-5), 572.310 Da. Adduct ions of $Na^+$ and $K^+$ were also observed, with m/z of 595.594 and 614.575, respectively. In order to confirm the identity of the ion at 573.603, it was further fragmented resulting in the spectrum at (C). Fragmentation pattern of the ion at 575.603 confirmed the identity of BK-(1-5).

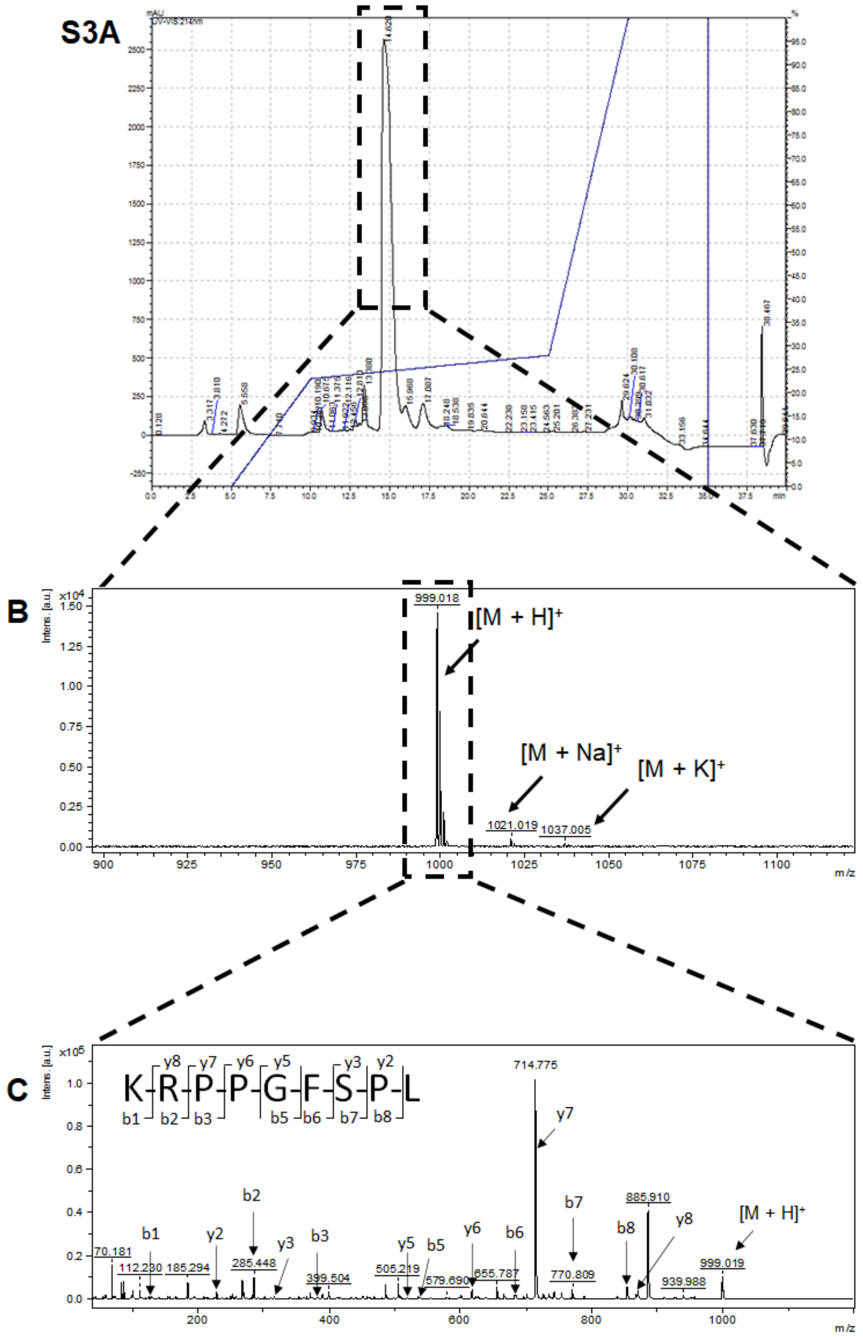


**Supplementary Figure S3 – Purification and characterization of Lys-(des-Arg$^9$,Leu$^8$)-BK-(1-9) synthesis products.** (A) Representative chromatogram from the purification process of crude Lys-(des-Arg$^9$,Leu$^8$)-BK-(1-9). The main peak at 14.8 minutes (in the black dashed box) was collected. (B) MALDI TOF/TOF mass spectrum of the collected fraction in the chromatogram in (A). A prominent ion in 999.018 $[M + H]^+$, with an experimental mass of 998.011 Da, corresponded to the theorical mass of Lys-(des-Arg$^9$,Leu$^8$)-BK-(1-9), 998.18 Da. Adduct ions of $Na^+$ and $K^+$ were also observed, with m/z of 1021.019 and 1037.005, respectively. In order to confirm the identity of the ion at 999.018, it was further fragmented resulting in the spectrum at (C). Fragmentation pattern of the ion at 999.018 confirmed the identity of Lys-(des-Arg$^9$,Leu$^8$)-BK-(1-9).

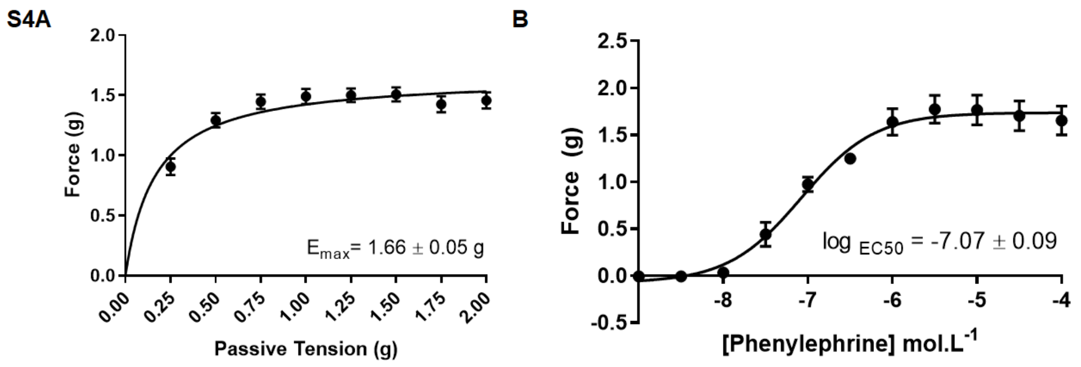


**Supplementary Figure S4 – Standardization of passive tension and concentration of phenylephrine in the vascular reactivity protocol.** (A) Passive tension curve on rat thoracic aorta rings, in which maximum contraction in each tension was elicited by KCl 6 x $10^{-2}$ $mol.L^{-1}$. Tension was raised by increments of 0.25g ranging 0.25 to 2g. Nonlinear regression of points (n=5). $E_{max}$ = 1.66 ± 0.05 g. Result expressed as mean ± SEM. (B) Concentration-response curve of phenylephrine on rat thoracic aorta rings. Nonlinear regression of points (n=5). $Log_{EC50}$ = 7.07 ± 0.09. Results expressed as mean ± SEM.

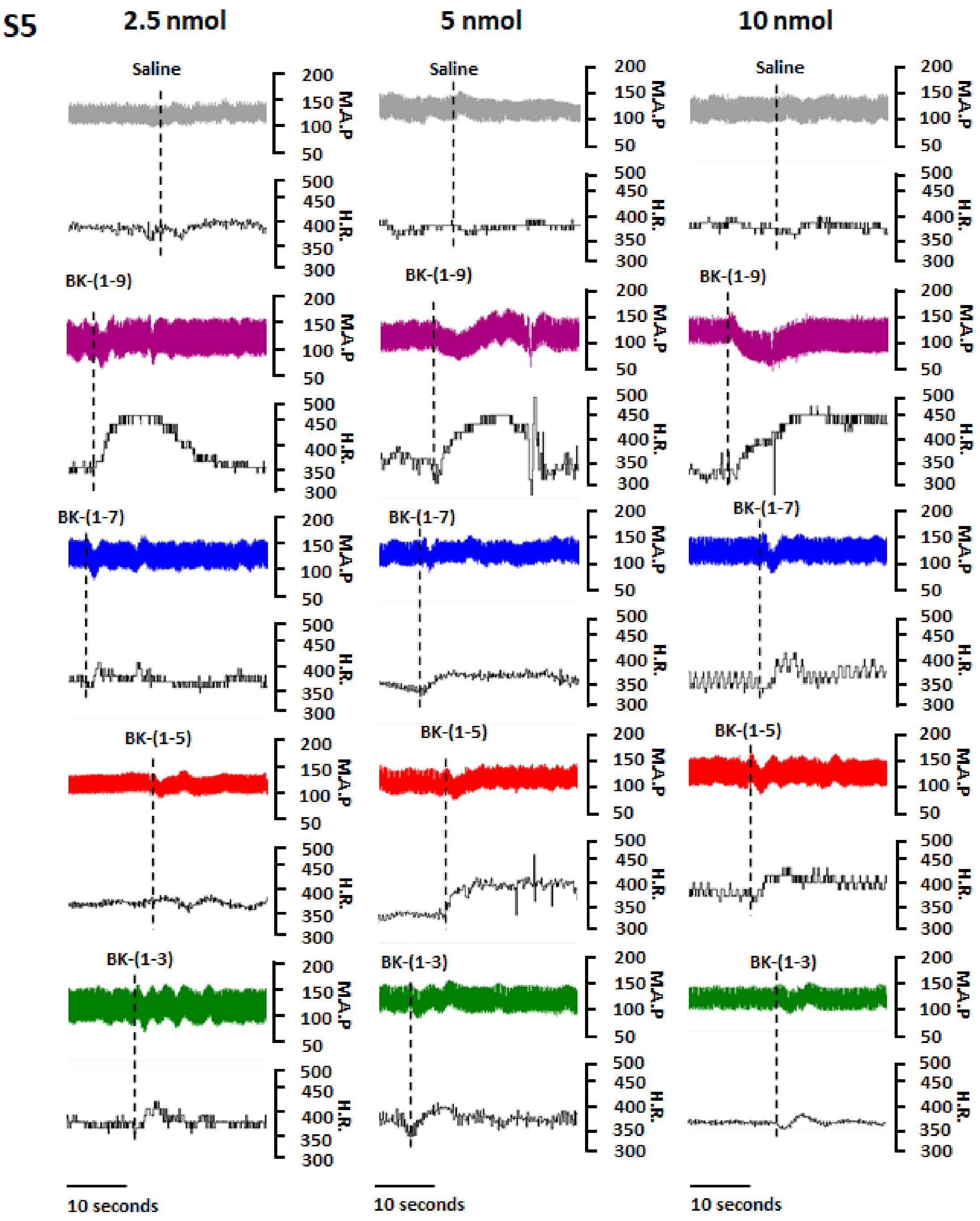


**Supplementary Figure S5 – Trace curves of dose-response curve hypotension mediated by BK-(1-9) and its fragments.** Changes of the mean arterial pressure (M.A.P.) and heart rate (H.R.) mediated by an intravenously (*i.v.*) administration of 0.1mL of saline or 0,1mL of BK-(1-9), BK-(1-7), BK-(1-5) or BK-(1-3) containing 2.5, 5 or 10nmol. Injections are depicted in dashed lines.

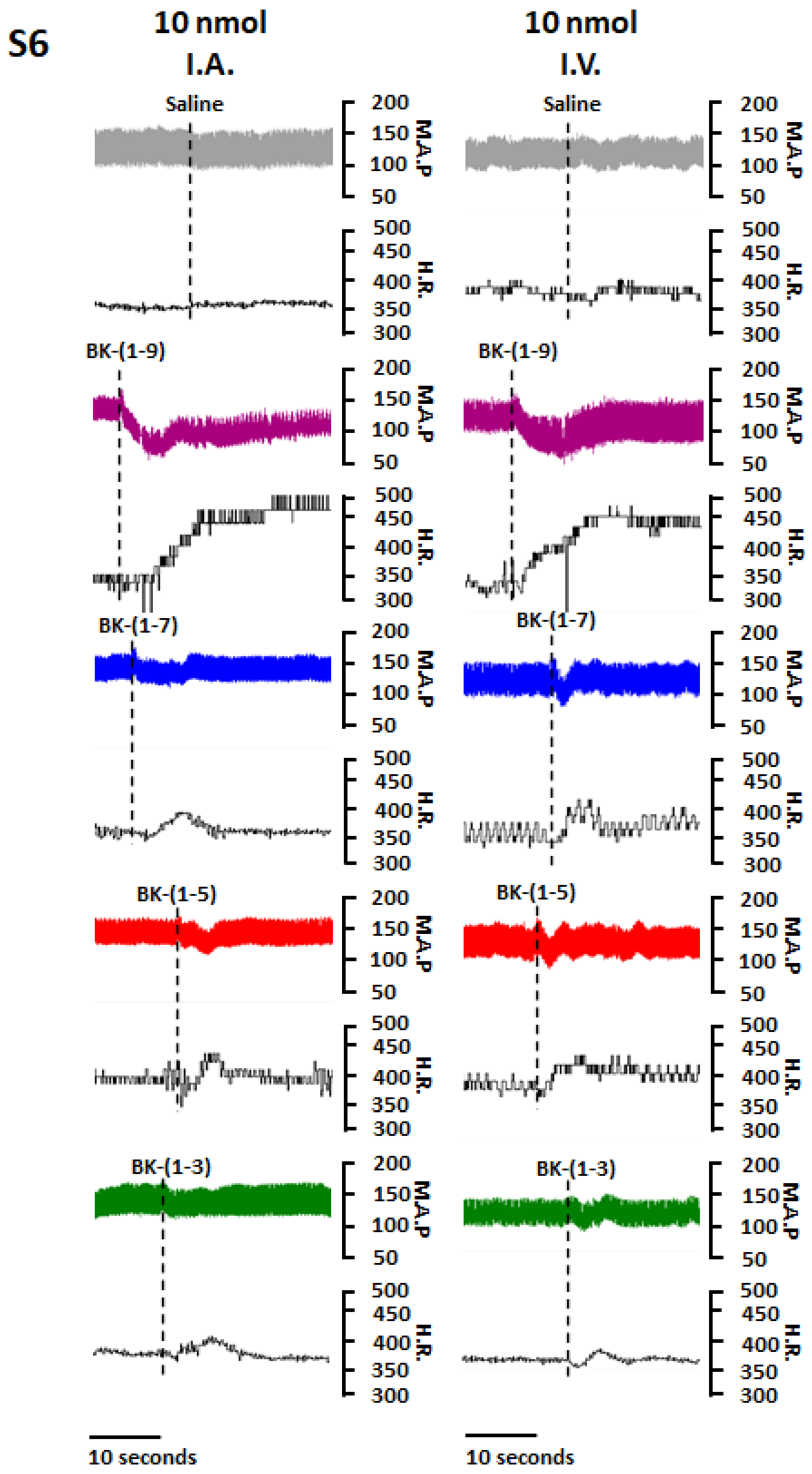


**Supplementary Figure S6 – Trace curves of intravenous or intraarterial administration on hypotension mediated by BK-(1-9) and its fragments.** Changes of the mean arterial pressure (M.A.P.) and heart rate (H.R.) mediated by an intravenously (*i.v.*) or intra-arterial (*i.a.*)

administration of 0.1mL of saline or 0.1mL of BK-(1-9), BK-(1-7), BK-(1-5) or BK-(1-3) containing 10nmol. Injections are depicted in dashed lines.

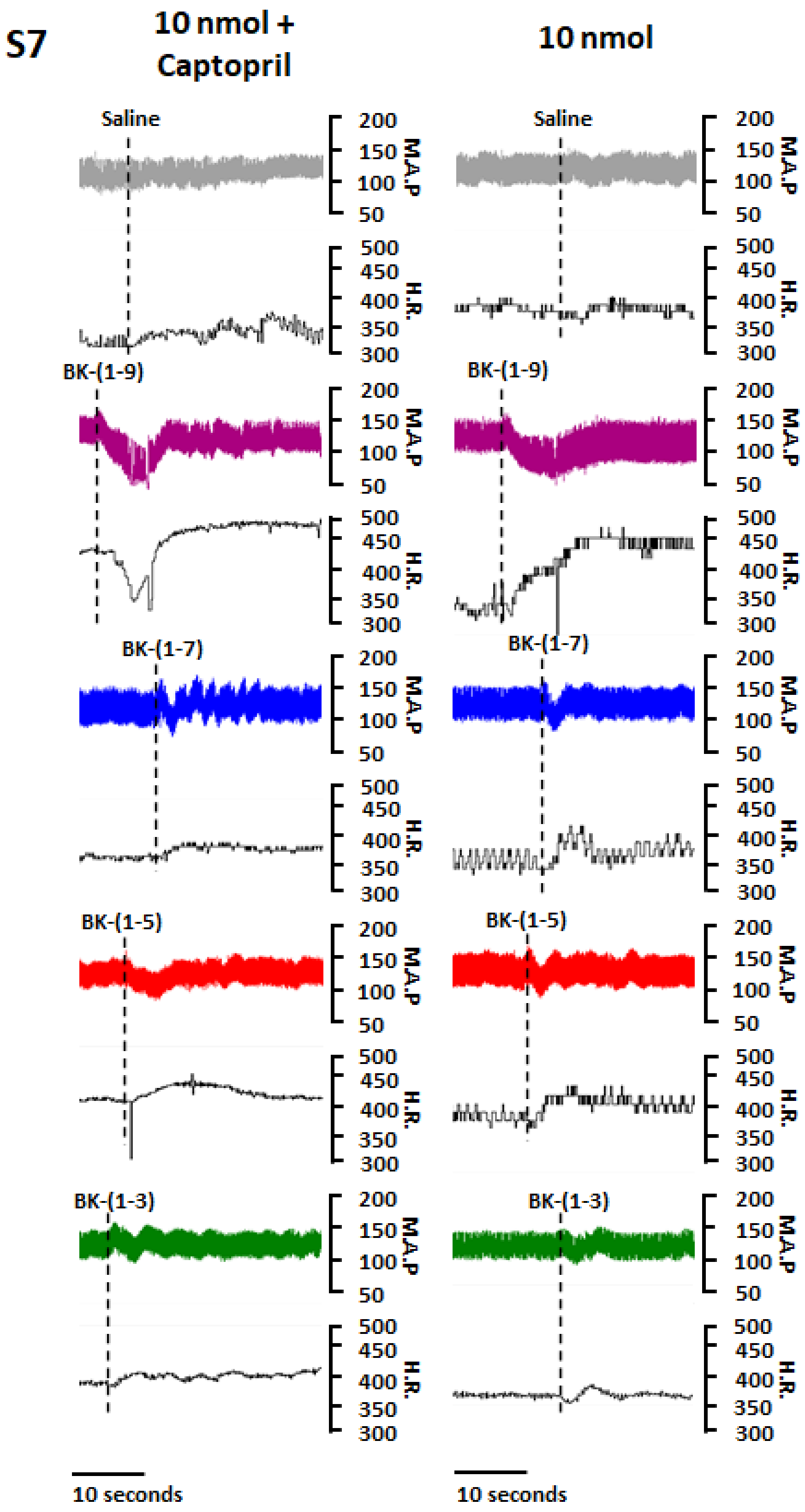


**Supplementary Figure S7 – Trace curves of intravenous or intra-arterial administration on hypotension mediated by BK-(1-9) and its fragments.** Changes of the mean arterial pressure (M.A.P.) and heart rate (H.R.) mediated by an intravenously (*i.v.*) administration of

0.1mL of saline or 0.1mL of BK-(1-9), BK-(1-7), BK-(1-5) or BK-(1-3) containing 10nmol. Captopril (5mg.kg$^{-1}$) was administered intravenously (*i.v.*) 20 minutes before administration of saline, BK-(1-9), BK-(1-7), BK-(1-5) and BK-(1-3). Injections are depicted in dashed lines.

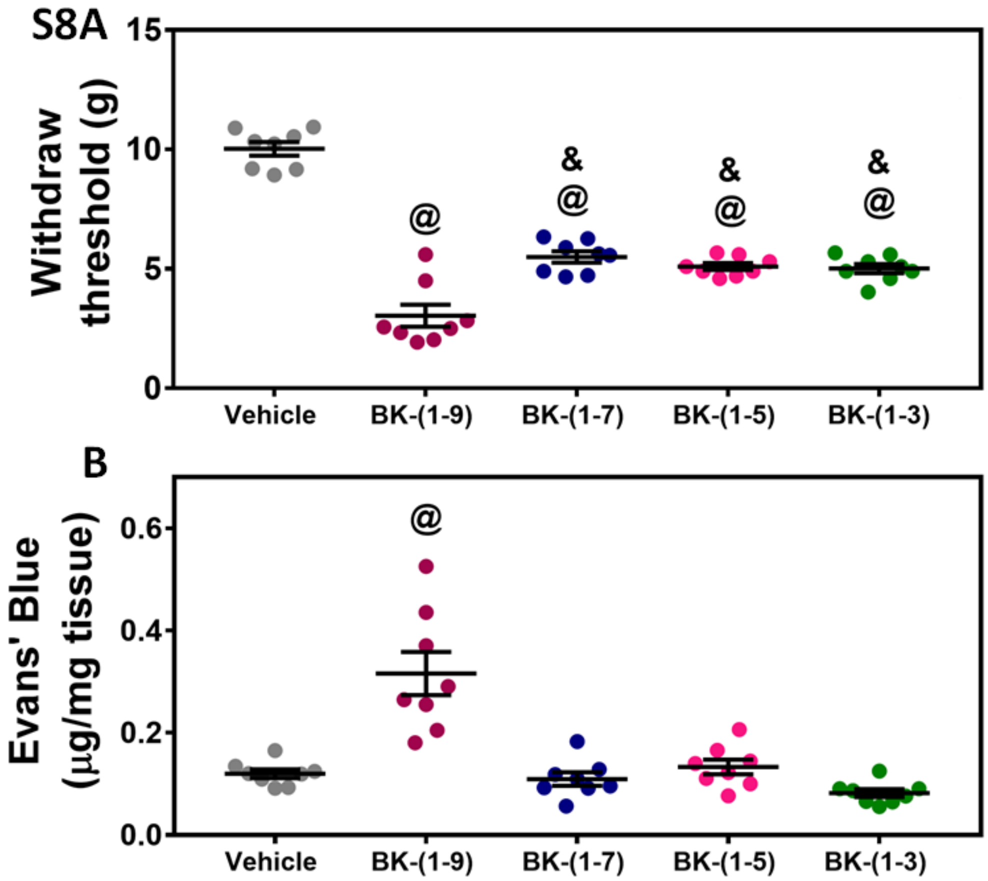


**Supplementary Figure S8 – *In vivo* proinflammatory actions of BK-(1-9) and its fragments.** (A) BK-(1-9), BK-(1-7), BK-(1-5) and BK-(1-3) increased the nociceptive reflexes when $10^{-8}$ mol in 10µL was administered intradermally into the footpad of mice of 8-12 weeks old. BK-(1-9) effect was twice as more potent than BK-(1-7), BK-(1-5) and BK-(1-3). Saline (vehicle) was used as control. One-way ANOVA with Tukey's multiple comparison *post-hoc* test. @ $p < 0.001$ when compared to control, & $p < 0.0001$ when compared to BK-(1-9) (n = 7). (B) BK-(1-9) induced plasma extravasation when $10^{-8}$ mol in 10µL was administered intradermally into the footpad of mice of 8-12 weeks old. BK-(1-7), BK-(1-5) and BK-(1-3) did not increase vascular permeability when administered at the same dose. Saline (vehicle) was used as control. One-way ANOVA with Tukey's multiple comparison *post-hoc* test. @ $p < 0.0001$ when compared to control (n = 7). All results are shown as mean ± SEM.

### SUPPLEMENTARY METHODS

#### Peptide Synthesis

BK-(1-9), BK-(1-5) and Lys-(des-Arg$^9$,Leu$^8$)-BK-(1-9) synthesized by solid-phase peptide synthesis, by the Fmoc strategy (for more details, please go to ). Briefly, Fmoc-Arg(Pmc)-NovaSyn-TGA, Fmoc-Phe-NovaSyn-TGA and Fmoc-Leu-NovaSyn-TGA (Merck) were used as starting material for the synthesis of BK-(1-9), BK-(1-5) and Lys-(des-Arg$^9$,Leu$^8$)-BK-(1-9), respectively. Elongation of the peptide chain was done by coupling the respective amino acid with 5 equivalents of Fmoc-amino acid, 5 equivalents of HBOt (Sigma-Aldrich), 4.9 equivalents of HBTU (Sigma-Aldrich) and 6 equivalents of DIPEA (Sigma-Aldrich) for one hour under constant stirring. Cleavage of crude peptide from the peptide-resin was made possible by a solution of TFA/triisopropylsilane/water at a proportion of 95:2.5:2.5% (v/v) for one hour under constant stirring. Crude peptides were stored at -20°C until usage.

#### Purification of crude products

Crude peptides were purified by high performance liquid chromatography (HPLC), in a Shimadzu CBM-20A system, coupled with two LC-20AT pumps and a UV-Vis SPD-20A detector. Mobile phase A consisted in a 0.1 % trifluoroacetic acid (TFA) (Sigma-Aldrich) solution in ultrapure type I water. Mobile phase B consisted in a 0.1% TFA solution in acetonitrile chromatographic grade (Sigma-Aldrich). Crude peptides were purified in a Discovery Bio Wide Pore C18, 10µm particle size, L x I.D 25cm x 10mm, in a 5 ml.minute$^{-1}$ flow. Purification of BK-(1-9): i) 0 to 5 minutes, 0% mobile phase B; ii) 5 ato10 minutes, 20% mobile phase B; iii) 10 to 25 minutes, 23% mobile phase B; iv) 25 to 30 minutes, 100% mobile phase B; v) 30 to 35 minutes, 100% mobile phase B; vi) 35 to 40 minutes, 0% mobile phase B. Purification of BK-(1-5): i) 0 to 5 minutes, 0% mobile phase B; ii) 5 to 15 minutes, 15% mobile phase B; iii) 15 to 25 minutes, 17% mobile phase B; iv) 25 to 30 minutes, 100% mobile phase B; v) 30 to 35 minutes, 100% mobile phase B; vi) 35 to 40 minutes, 0% mobile phase B. Purification of Lys-(des-Arg$^9$,Leu$^8$)-BK-(1-9): i) 0 to 5 minutes, 0% mobile phase B; ii) 5 to 10 minutes, 23% mobile phase B; iii) 10 to 25 minutes, 28% mobile phase B; iv) 25 to 30 minutes, 100% mobile phase B; v)

30 to 35 minutes, 100% mobile phase B; vi) 35 to 40 minutes, 0% mobile phase B. Most abundant peaks of each chromatogram were collected for further characterization. Fractions were lyophilized and store at -20°C until usage.

**Characterization by mass spectrometry**

All characterization procedures were done in a MALDI TOF/TOF Autoflex III Smartbeam mass spectrometer. Purified peptides were resuspended in 0.1% TFA solution in ultrapure water type I and then homogenized in a 1:1 proportion with α-cyano-4-hydroxycinnamic acid matrix (Sigma-Aldrich). This mixture was transferred to a MTP 384 Polished Steel plate (Bruker Daltonics) and left to dry. For calibration purposes, a Peptide Calibration Standard II (Bruker Daltonics) was homogenized in a 1:1 proportion with α-cyano-4-hydroxycinnamic acid matrix and left to dry. Before spectra acquisition, an external calibration was performed. Peptide mass spectra was acquired in positive reflected mode with 300 laser shots, while acceleration voltage was held between 15-20kV. Tandem mass spectra were collected by selecting the major m/z for fragmentation using the LIFT mode.

**Standardization of passive tension**

Rat thoracic aortic rings were carefully coupled to a force transducer and left to stabilize for 60 minutes at the starting passive tension of 0.25g. Passed the stabilization period, maximum contraction was elicited by KCl at 6 x $10^{-2}$ $mol.L^{-1}$. After contraction, the rings were thoroughly washed with fresh Krebs-Henseleit solution until tension was reestablished. Passive tension was increased in increments of 0.25g until 2g. Maximum contraction was only performed after stabilization of a new passive tension.

**Standardization of concentration of phenylephrine**

Rat thoracic aortic rings were carefully coupled to a force transducer and lefr to stabilize for 60 minutes at the optimal passive tension of 1.60g. After the stabilization period, a concentration-response curve of phenylephrine was performed, where the tested concentrations ranged from $10^{-9}$ to $10^{-4}$ $mol.L^{-1}$. Results were submitted to a nonlinear fit curve to obtain $EC_{50}$, which is the concentration used for suboptimal aortic ring contraction.

### *In vivo* plasma extravasation experiment

Plasma extravasation was performed in 8-12 weeks old C57Bl/6 mice as previously reported (48). Briefly, 10 µL of $10^{-8}$ mol of BK-(1-9), BK-(1-7), BK-(1-5) or BK-(1-3) was injected intradermally in the hind paw and sterile saline was injected in the same volume on the contralateral paw as control. Immediately afterwards, Evans blue (Merck) solution at 2% was injected intraorbitally for measurement of plasma extravasation. After 30 minutes of peptide administration, mice were euthanized by cervical dislocation and footpads were dissected. The footpads were dried at 60°C for 24 hours and the tissue was weighted. Evans blue was extracted by formamide for 72 hours under constant stirring and then the absorbance of this final solution was measured by a spectrophotometer at $\lambda$ = 620nm. Results were expressed as µg of Evans Blue per mg of dried tissue.

### *In vivo* nociceptive assay

Potential nociceptive reflexes mediated by BK-(1-9), BK-(1-7), BK-(1-5) and BK-(1-3) were evaluated as previously described (47). 8-12 weeks old C57Bl/6 mice were submitted to 10 µL intradermal hind paw injection of BK-(1-9), BK-(1-7), BK-(1-5) or BK (1-3) at a dose of $10^{-8}$ mol. Saline was injected on the contralateral paw as control in the same volume. Animals were placed individually in acrylic cages with wired floor and submitted to an adaptation period of 20 minutes. Evaluation of mechanical nociceptive reflex was assessed with a pressure-meter (Insight Instruments) by applying a crescent pressure in the hind paw until the mice evoked a dorsiflexion withdrawn reflex. The pressure by which this event occurred, expressed in grams (g) was registered.
